# Occurrences of protonated base triples in RNA are determined by their cooperative binding energies and specific functional requirements

**DOI:** 10.1101/2021.01.10.426061

**Authors:** Antarip Halder, Ayush Jhunjhunwala, Dhananjay Bhattacharyya, Abhijit Mitra

## Abstract

With wide ranging diversity in their geometries, binding strengths and chemical properties, noncanonical base pairs are equipped to intricately regulate and control the structural dynamics of RNA molecules. Protonation of nucleobases adds to the diversity. Compared to the unprotonated scenario, on one hand they open up new alternatives for base pairing interactions (*Class I*) while on the other, they modulate the geometry and stability of existing base pairing interactions (*Class II*). In both cases, compensation of the energetic cost associated with nucleobase protonation at physiological pH, can be understood in terms of protonation induced restructuring of charge distribution. This not only leads to modifications in existing base-base interactions but often also leads to additional stabilizing interactions, resulting in the formation of protonated base triples. Here we report our detailed quantum chemical studies, in conjunction with structural bioinformatics based analysis of RNA crystal and NMR structure datasets, probing into the contribution of such protonated triples in the structural dynamics of RNA. Our studies revealed more than 55 varieties of protonated triples in RNA, some of which occur recurrently within conserved structural motifs present in rRNAs, tRNAs and in other synthetic RNAs. Our studies suggest that high occurrence frequencies are associated with protonated triples which satisfy the specific structural requirements of conserved motifs where they occur. For example, protonated triples with flexible geometries are involved in the formation of tertiary contacts between different distant motifs. Stabilization of protonated base pairs, through the induction of additional energetically cooperative interactions, appears to be another factor. These results provide significant insights into the sequence-structure-function relationships in RNA.

## Introduction

RNAs are essential for a wide variety of cellular processes.^1,2^ It has long since been known^3^ that rRNA, tRNA and mature mRNA constitute the key components of the protein synthesis machinery in both prokaryotes and eukaryotes.^4^ However, recent research has revealed a plethora of non(protein)coding RNAs.^5^ In eukaryotic cells, their roles include repression of translation by miRNAs,^6^ inhibition of transcription by siRNAs,^7^ splicing of pre-mRNA by snRNAs,^8^ post-transcriptional modifications in nascent rRNAs by snoRNAs,^8^ tRNA processing by RNase-P,^9^ etc. In addition to these, regulation of gene expression by riboswitches and catalysis of biophysical processes by ribozymes are common in prokaryotes.^10^

The polyanionic RNA molecule needs to fold into its active native structure to perform its function.^11^ It was proposed that metal ions present in the cellular environment act as counterions neutralizing the negative charges of the backbone and cause the initial collapse of the RNA chain.^12–14^ The collapsed RNA further weaves an intricate net of secondary and tertiary interactions to stabilize the final folded functional structure. Extensive scientific investigations notwithstanding, a detailed understanding of how RNAs can play such a wide variety of structural and functional roles using only four chemically similar nucleotides and their modification, continue to elude us.^15^

The RNA polynucleotide is formed through phosphodiester linkages between adjacent nucleotides involving the 3′-OH of the i^*th*^ nucleotide with the 5′-phosphate of the (i-1)^*th*^ nucleotide. Each nucleobase can be characterized by their three edges (Watson-Crick (W), Hoogsteen (H) and sugar (S)) and each edge has a unique base specific set of hydrogen bond donor and acceptor atoms. ^16^ It is now well accepted that pairs of nucleobases in RNA form hydrogen bonded planar base pairs with each other in different interacting geometries.^17^ Leontis and Westhof categorized these base pairing geometries into 12 classes based on the interacting edges of the two bases (WW, WH, HH, WS, HS and SS) and mutual orientation of the glycosidic bonds (cis or trans).^18^ So with four types of naturally occurring nucleobases there exits a total of 156 distinct base pairing possibilities. Details of the counting method and nomenclature used to annotate different base pairs are discussed in Supporting Information. Some of these possibilities lack the complementary hydrogen bonding acceptor-donor network between the interacting edges while some are ruled out due to steric constraints. A recent study indicates that about 118 different base pairs are actually observed in RNA crystal structures.^19^ All these base pairs, apart from the most commonly occurring canonical A:U W:W Cis and G:C W:W Cis base pairs, are called noncanonical base pairs. Detailed quantum chemical studies have revealed that noncanonical base pairs display a rich variation in geometry, stability, and physicochemical characteristics.^20–26^ Further, analyses of their occurrence contexts in RNA crystal structures suggest that many of these noncanonical base pairs are associated with specific motifs or structural contexts. For example, G:A S:H Trans forms the loop closing pair^27^ in all instances of the ubiquitous,^28^ and exceptionally thermostable,^29^ GNRA tetraloop motif. It may therefore be hypothesized that noncanonical base pairs, given their wide ranging physicochemical characteristics, play significant roles in shaping the structures and in controlling the dynamics of individual motifs and hence, of the overall RNA.^30^

RNA nucleobases often undergo chemical modifications, which add to the diversity in the repertoire of noncanonical base pairs in RNA. These include post-transcriptional modification,^31^ metal ion binding,^32^ tautomerizatio,^33^ ionization,^34^ etc. Detailed studies have been carried out to understand the influence of post-transcriptional modifications^35,36^ and metal ion binding^37^ on the geometry and stability of RNA base pairs. It is important to note that, some of these modifications (protonation, tautomerization, etc.) enable the nucleobases to participate in newer kinds of base pairing interactions, which are different from those observed with corresponding unmodified bases. Involvement of rare nucleobase tautomers in causing mutagenic mispairings during DNA replication is well known.^38,39^ It has been suggested that different putative minor tautomeric forms of nucleobases may enhance their structural and functional diversity.^40^ Nevertheless, reported examples of RNA base pairing interactions suggestive of the involvement of rare tautomers of nucleobases are very few.^41^ In contrast, examples of protonated base pairs are abundant in RNA crystal structures.^42^ Several base-base contacts have been noted in the crystal structures of RNA where two hydrogen bonding acceptor atoms (i.e., electronegative imino-Nitrogen or carbonyl Oxygen atoms) are closely spaced. This can only happen if one of them carries an extra proton, making the corresponding base positively charged, thus changing the electrostatic repulsion to attractive hydrogen bonded pairs. Such instances of nucleobase protonation are categorized as *Class I* protonation. In a nonredundant set of RNA crystal structures, 18 such distinct types of such base pairing interactions have been observed.^42^ Diverse roles of Class I protonated base pairs are observed in RNA which include modulation of the maturation of oncogenic microRNA-21,^43^ stabilization of the tertiary structure of GTP-binding RNA aptamer,^44^ controlling the conformational change between the two steps of splicing in a self-splicing bacterial Group II intron,^45^ maintaining encapsidation efficiency of turnip yellow mosaic virus,^46^ etc. They have been found to stabilize important structural motifs also in DNA, e.g., i-motif,^47,48^ triple helical DNA,^49,50^ etc. When the loaded proton is not directly involved in base pairing (*Class II* protonation), it can participate in catalytic processes with roles similar to those of charged amino acids involved in the catalytic actions of proteins.^51,52^ Circumstantial evidences of *Class II* protonated bases pairing with other neutral bases via one of their unprotonated edges have been reported by computational^53,54^ and X-ray crystallographic^55^ studies. For example, N1 protonation of adenine is implicated in alternate A:A H:H Trans pairs that form the parallel double helix in a poly(rA) staggered zipper.^55^ *Class II* protonation significantly modify the geometry and stability of those noncanonical base pairs.^53,54^ Interestingly, nucleobase protonations remain ‘invisible’ to the conventional biophysical methods, due to absence of hydrogen atom coordinates in the X-ray crystal structures and to the ambiguity in their assignment in NMR structures. So theoretical and computational techniques constitute an important approach complementing experiments towards understanding the role of nucleobase protonation in RNA structural dynamics.

The pK_*a*_ values of nucleobases are 2-3 units away from biological pH (≈7.4).^56,57^ So under normal physiological conditions, nucleobase protonation is energetically unfavorable. The general perception is that the energetic cost of protonation is compensated by the overall stabilization achieved due to involvement of the protonated base in base pairing and other inter-nucleotide interactions.^34,42^ It is important to characterize such inter-nucleotide interactions for a comprehensive understanding of the contribution of nucleobase protonations in the structural dynamics of RNA. Both experimental^58,59^ and computational^42,60^ studies show that interaction energies of noncanonical base pairs with *Class I* type protonation are usually higher than those of canonical A:U and G:C pairs. This also holds for base pairs with *Calss II* protonation.^53,54,61^ Interestingly, *Class I* protonated base pairs may further be involved in base triple formation by interacting with another neutral base.^42,60^ In these base triples, the third base can interact with either the neutral base or with the protonated base of the corresponding protonated base pair. These two situations are illustrated in Figure 1. The former case has been discussed earlier in the context of Triplex Forming Oligonucleotides (TFO),^62^ which are known for their biological activities and therapeutic applications.^63^ It has also been shown that hydrogen bonding potential of the free edges of the neutral base is enhanced in proton mediated base pairs, which, in turn, is expected to facilitate protonated base triple formation.^60^ On the other hand, the latter situation is particularly interesting because the protonated base becomes the central base which forms a *Class I* protonated base pair with one of the terminal bases, while engaging in *Class II* type protonated base pairing with the other. Hence, protonated base triples constitute an interesting set of systems for studying how the consequences of *Class I* and *Class II* type nucleobase protonations cooperate with each other.

**Figure 1:**
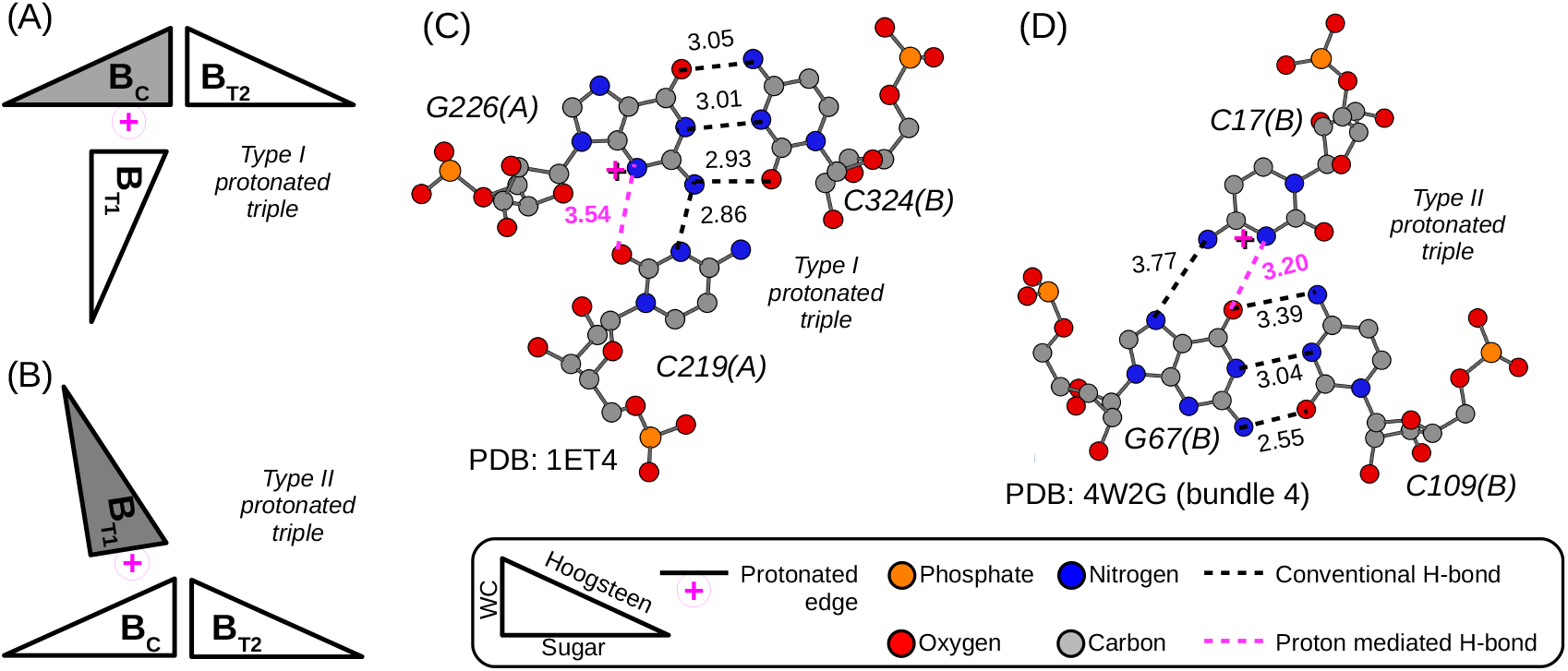
A protonated base triple is composed of a central base (B_*C*_) and two terminal bases (B_*T*1_, B_*T*2_), where B_*C*_ forms two different base pairs with B_*T*1_ and B_*T*2_ and one of these two pairs is a *Class I* protonated base pair. We have categorized the protonated triples into two types. In Type I, B_*C*_ is protonated, whereas in Type II either B_*T*1_ or B_*T*2_ is protonated. Schematic representations of the base-base interaction patterns (as per LW scheme ^18^) of Type I and Type II protonated base triples are shown in panel (A) and (B), respectively. The protonated and neutral bases are represented as filled and empty triangles, respectively. One example crystal structure of (C) Type I and (D) Type II base triples are also shown. The inter base hydrogen bonds are shown in broken lines with donor-acceptor distances reported in Å. In this figure and other figures of this paper, the proton mediated hydrogen bond is shown in magenta color.

In this work the RNA crystal structure datasets are analyzed to study the occurrence contexts of different protonated base triples, followed by quantum mechanical evaluation of their intrinsic structure and stability. The results provide some rationale behind abundance (or lack of it) of certain types of protonated triples in RNA structure data sets. They also explain how the intrinsic geometry and stability of different protonated triples determine their belongingness to specific RNA structural motifs.

## Methods

### Annotation of Type I and Type II protonated base triples

A base triple is composed of three nucleobases (say, B_*T*1_, B_*C*_ and B_*T*2_) where the central base B_*C*_ forms two base pairs with the two terminal bases B_*T*1_ and B_*T*2_, respectively. If any of these two base pairs (B_*C*_:B_*T*1_ or B_*C*_:B_*T*2_) is a *Class I* protonated pair, then the resulting triple is identified as a protonated base triple. There could be two major varieties of the composition of a protonated base triple. Either the protonated base occurs at the central position (B_*C*_), or at one of the terminal positions (B_*T*1_/B_*T*2_). We categorize the former case as Type I (Figure 1A) and the later as Type II (Figure 1B). There could be a third variety where the triple is cyclic, i.e., the two terminal bases are paired with each other. However, we could not find any example of cyclic protonated triples in the RNA structural datasets studied in this work. In principle, there could be another class of protonated triples, where either (a) any of the terminal bases gets protonated at any of their free edges and subsequently form *Class II* protonated base pair with the central base, or (b) the central base may get protonated at one of its free edges and form *Class II* protonated base pairs with both the terminal bases. However, analyzing the occurrence contexts and energetics of this class of protonated triples are beyond the scope of this study as conventional base triple detection algorithms like BPFIND,^64^ 3DNA,^65^ etc. can not identify base triples having only *Class II* protonation and no *Class I* protonation.

To simplify the systematic analysis, each triple has been annotated by the base combination and interaction geometries of the two constituent base pairs. While representing the base combinations, the terminal bases are kept within parenthesis and the central base is kept outside of it, e.g., B_*C*_(B_*T*1_B_*T*2_). Representation of interaction geometries contain B_*C*_B_*T*1_ and B_*C*_B_*T*2_ base pairing geometries separated by a ‘/’. For the protonated base, ‘+’ represents the protonated Watson-Crick edge and ‘z’ represents the protonated Sugar edge. Note that, since purines can not form base pairs with their protonated Hoogsteen edge^60^ and pyrimidines do not have a protonable site at their respective Hoogsteen edges (C-H edges), no extra symbol is required to designate the protonated Hoogsteen edge. Hence, the two protonated base triples illustrated in panel A and B of Figure 1 can be annotated as G(CC) z:WC/W:WC and G(CC) H:+T/W:WC, respectively. The same protocol is used to annotate bifurcated triples where the two terminal bases interact with the same edge of the central base. For bifurcated base triples of Type I, two terminal bases interact with the protonated edge of the central base. Usually in those cases only one base pair, out of the two, involves the proton mediated inter-base hydrogen bond. In such cases, special symbols like ‘+’ or ‘z’ are used to repesent the interaction geometry in one base pair and regular symbols like ‘W’ and ‘S’ are used in the other.

### Context analysis

Detailed analysis, of the respective structural contexts, have been carried out for different triples, in order to determine their putative association with distinct structural motifs. For this purpose, two dimensional maps of the secondary structures of the different ribosomal RNAs provided by the RiboVision^66^ server are used as a template. To understand the effect of these proton mediated tertiary contacts on the folding of RNA chains, structural alignment of relevant RNA chains are carried out using the SETTER web server.^67^ For further visual inspection, the RNA fragments containing protonated base triples are rendered using molecular visualization tools like VMD and Jmol.^68^

### Choice of RNA structure datasets

RNA molecules often display large conformational variations, and hence variations in their inter-nucleotide interaction patterns, in response to changes in the local environment as well as to even small sequence variations.^69,70^ Hence, a conventional nonredundant dataset is not expected to provide a comprehensive repertoire of different instances of base triples present in RNA. On the other hand, the statistical biases inherent in redundant datasets are expected to skew their respective occurrence frequency data, leading to misleading inferences regarding their significance. In order to address these and related reliability issues, we have used three datasets for our study. All the RNA X-ray crystal structures with a resolution better than 3.5 Å (total 1873 structures) and all the solution NMR structures containing RNA fragments (total 591 structures), which were available on PDB server on September, 2015 constitute a curated dataset, henceforth to be referred to as the ‘large dataset’. In addition to this, we consider two non-redundant sets of RNA crystal structures. One is provided by the version 1.89 of the Nucleic Acid Database (NDB)^71^ and the other is provided by the HD-RNAS.^72^ The ‘NDB dataset’ consists of 838 structures shortlisted after applying a resolution cut-off of 3.5 Å on the structures available in the NDB database. The ‘HD-RNAS dataset’, on the other hand, consists of 167 RNA crystal structures, which have been shortlisted by applying a resolution cut-off of 3.5 Å and length cut-off of 30 nucleotides in order to exclude the small synthetic RNA constructs. It may be mentioned here that filters based on chain length, R-factor, resolution and sequence similarity, etc., have already been applied while including the representative structures present in HD-RNAS. The NDB and HD-RNAS are not subsets of the ‘large dataset’ since both the nonredundant datasets contain some structures which were marked obsolete and were removed from the distribution of released PDB entries before curation of the ‘large dataset’. PDB Ids of all the crystal and solution NMR structures of the three datasets are respectively listed in Supporting Information. The same sets of RNA structures were used in earlier works^37,73,74^ and the statistical trends observed in them collectively provided comprehensive insights.

### Modeling base triples for quantum mechanics (QM) based calculations

BPFind software^64^ has been used to detect base triples involving *Class I* protonated base pairs in all the three datasets mentioned above. We have developed an in-house code to identify whether the detected protonated triple is of Type I or of Type II. Occurrence frequencies of the protonated triples in these datasets are reported in Supporting Information (Table S1, S2 and S3). Within the crystal environment the geometries of the hydrogen bonds, which mediate the RNA base pairs, usually deviate from their ideal geometry due to environmental factors. BPFind reports a composite parameter called ‘E-value’, based on the geometrical variables of the hydrogen bonds involved, that quantifies the ‘goodness’ of the resultant base pair. The lower the E-values the closer are the base pairs to their corresponding predicted ideal hydrogen bonding geometries. Building upon this concept, the ‘goodness’ of base triples were quantified in terms of the sum of the E-values of the two constituent base pairs. Out of all the detected instances belonging to any given type of protonated base triple (say, G(CC) H:+T/W:WC), the instance with the least E-value sum was considered to be closest to the ideal planar geometry. Those examples are therefore selected for subsequent QM calculations (Table S4).

The initial geometries of these interacting systems have been modeled by adding hydrogen atoms at appropriate positions and by replacing the non-participating sugar moieties by methyl groups. The sugar moieties, with O5′ replaced by H and O3′ valency satisfied using a methyl group, were however retained in the models, whenever the sugar edge of a base was involved in the pairing interactions. Such a strategy has been shown to reduce the computational cost, without affecting the essential chemical interactions involved.^75^ GaussView^76^ package has been used to perform the modeling and editing of the initial structures.

### Geometry optimization

The modeled triples were geometry optimized in gas phase using the hybrid meta-GGA functional M05-2X^77^ and Pople type^78^ split valance double-ζ basis set 6-31++G(2d,2p) which is augmented with (a) two sets of a d-type polarization function for all nonhydrogen atoms, and (b) two sets of p-type polarization function for hydrogen atoms, also including s-p diffused orbitals for nonhydrogen atoms. M05-2X functional can predict the equilibrium geometry^79^ and interaction energy^80^ of small bimolecular van der Waals complexes with remarkable accuracy. As suggested in earlier publications by our group^37,53,54,81^ and others,^82–84^ its performance is also reliable for noncovalently bonded molecules of biological relevance, especially with nucleobases.

Nevertheless, we have also benchmarked its performance for calculating the geometries of protonated triples with five other DFT functionals which have frequently been used in literature for studying DNA/RNA base pairs and related systems. Performance of M05-2X was consistent with these five hybrid-GGA DFT functionals, viz. B3LYP,^85–87^ CAM-B3LYP,^88^ B3LYP-D3,^89,90^ PBE0^91^ and ωB97-XD^92^ (Table S5). We selected one Type I (G(CC) z:WC/W:WC, Figure 1A) and one Type II (G(CC) H:+T/W:WC, Figure 1B) triples for this test. We compared the optimized geometries obtained using different DFT functionals on the basis of Root Mean Square Deviation (RMSD) between them. To calculate the RMSD between two structures, first they are superposed. We carried out two different protocols for superposition, (a) we superpose two optimized triples with respect to all non-hydrogen atoms and call the corresponding RMSD as RMSD(1) and (b) we superpose them with respect to only the central base and call the corresponding RMSD as RMSD(2). RMSD(1) quantifies the overall deviation and RMSD(2) helps in probing the deviation in individual base pairs. VMD software^93^ has been used for this purpose. We have further verified that the optimized geometries obtained using the 6-31++G(2d,2p) basis set are also consistent with the optimized geometries obtained using a larger basis set 6-311++G(2df,2pd) (Table S5). Note that it is a triple-ζ split-valance basis set augmented with (a) one set of a d-type and one set of a p-type polarization function for all nonhydrogen atoms, and (b) one set of a p-type and one set of a d-type polarization function for hydrogen atoms and includes s-p diffused orbitals for nonhydrogen atoms. All the quantum chemical calculations are performed using Gaussian 09 software. ^94^

We used natural population analysis (NPA) to estimate the partial charge distribution around the polar atoms of the bases. NPA is performed using NBO package^95^ implemented in Gaussian 09 software. ^94^ We have also performed the quantum theory of atoms in molecules (QTAIM) analysis^96,97^ on selected protonated triples using the AIMALL package.^98^

### Interaction energy calculation

Interaction energies of the M05-2X optimized geometries are calculated at MP2/aug-cc-pVDZ level. ^99^ For comparison we also have calculated the interaction energies using both M05-2X and ωB97XD functional with 6-311++G(2df,2pd) basis set. ωB97XD is based on Becke’s B97 exchange-correlation functional^100^ and includes both long range correction to the non-Coulomb part of exchange functional and Grimme’s second order explicit dispersion correction.^101,102^ Recent benchmarking of the performance of different DFT functionals for studying the optimized geometry, stability and cooperativity of DNA triplexes have shown that the B97-D functional (Becke’s functional with explicit dispersion correction) provides results consistent with higher levels of theory.^103^ Hence we included the ωB97XD functional for analysis.

The interaction energies are calculated as, ΔE = Electronic energy of the triple - Σ(Electronic energy of the individual bases). Interaction energies are further corrected for Basis Set Superposition Error (E^*BSSE*^) and deformation energy correction (E^*def*^). E^*BSSE*^ values have been calculated using counterpoise method^104^ considering the triple as a three body system. The deformation energy of a base triple formed by bases 1, 2 and 3 has been calculated as, 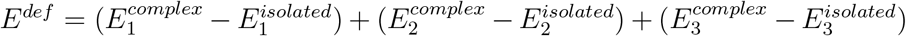 where 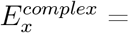 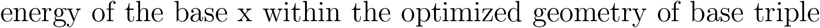 and 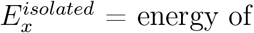 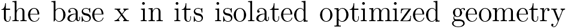. Hence the final interaction energy is, E^*int*^ = ΔE + E^*BSSE*^ + E^*def*^. However, due to computational constraints we can not calculate E^*BSSE*^ at MP2 level. Further, the Conductor-like Polarizable Continuum Model (CPCM)^105^ has been implemented to study the effect of the solvent screening on the electrostatic component of the interaction energies.

We calculate cooperative energies (CE) of the optimized base triples. Cooperative energy of a many body system is the difference between its interaction energy and the sum of the interaction energies of all its two body components. Therefore, the cooperative energy for a three body system (123) has been calculated as,

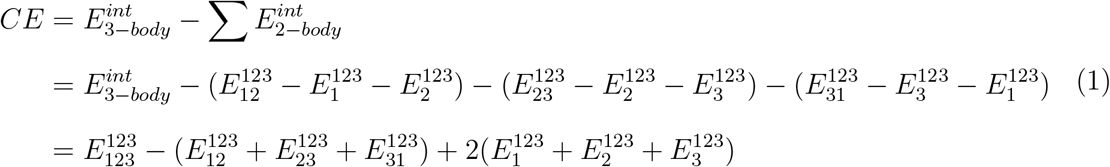

where, 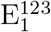 indicates the energy of the monomer 1 (in the optimized geometry of 123) calculated using the basis set of the complete 123 triple. Similarly, 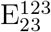 indicates the energy of the 23 pair (in the optimized geometry of 123) calculated using the basis set of the complete 123 triple.^106^ The CE of a base triple quantifies the influence of *Base-2* on the binding affinity of *Base-1* towards *Base-3*. Charge redistribution and geometric rearrangements resulting out of base pairing between *Base-1* and *Base-2* may improve the binding affinity of *Base-3* to *Base-1* and/or *Base-2*. In that case, the resulting base triple will have higher (more negative) interaction energy than the sum of interaction energies of its constituent base pairs. The resulting CE will be negative. Otherwise the resulting base triple will have CE > 0 and will be called as ‘anti-cooperative’. Due to computational constraints we carried out CE calculations at DFT level only. However, the calculations are performed using two DFT functionals M05-2X and ωB97XD with the larger basis set 6-311++G(2df,2pd).

## Results and Discussions

### Type II Protonated Base Triples Are More Frequently Observed Than Type I

We have identified the frequently occurring protonated triples in RNA crystal structures, analyzed their context of occurrences and carried out QM based analysis of their intrinsic geometry and electronic properties. Analysis of the two non-redundant datasets (HDRNAS and NDB) and the larger dataset suggest that Type II protonated base triples (i.e. base triples having a protonated base at one of the terminal positions) occur more frequently in RNA. The larger dataset contains 958 instances of protonated base triples, out of which 546 are of Type II. Similarly, 26 out of 36 instances of protonated base triples in HDRNAS and 35 out of 59 instances of protonated base triples in NDB are of Type II.

The 95 instances of protonated triples found in the two non-redundant datasets belong to 19 distinct types of base triples as characterized by (i) the identity of the three constituent bases and (ii) their geometry of interaction (Table 1). In addition to these 19 types of base triples, 39 other types of protonated triples are found in the larger dataset. However, the total number of occurrences of those 39 triples are insignificant (73 out of total 958 instances) and have not been studied in detail in this work. However, among those 39 base triples that occur exclusively in the larger dataset, the occurrence frequency of the G(GU) z:HT/W:WC triple is the maximum (6 instances) and therefore seeks special attention. Hence, along with the 19 base triples that occur in non-redundant datasets, the G(GU) z:HT/W:WC triple has also been shortlisted for detailed context analysis and QM calculations (Table 1). These 20 systems occur in different rRNAs, tRNAs and synthetic RNA constructs. Some of the triples also have recurrent occurrences in ribosomal RNAs (Table 1). Note that this set of 20 protonated base triples consists 10 Type I triples (Systems 1, 7, 9, 10, 12, 15, 17, 18, 19 & 20) and 10 Type II triples (Systems 2, 3, 4, 5, 6, 8, 11, 13, 14 & 16). This is a large enough sample set for studying contribution of protonated base triples in RNA’s structural dynamics. At the same time this constitutes an unbiased set for probing why Type II triples are favored over Type I.

**Table 1:** Frequency and context of occurrence of 19 base triples (with protonated bases) that are found in the two non-redundant datasets (HDRNAS and NDB). 9 of them have the protonated base at the center (Type I) and 10 of them have the protonated base at one of the terminal positions (Type II). System 20 belong to Type I category but does not occur in any of the non-redundant datasets. It is included due to its high occurrence frequency in the larger dataset. Out of these 20 pairs *System 11*, *15* and *19* are bifurcated base triples, where both the terminal bases interact with the same edge of the central base.

| Sys. No. | Base Triple | Category | Occurrence Frequency |  |  | Types of RNAs where the triples are found |
| --- | --- | --- | --- | --- | --- | --- |
|  |  |  | Larger Dataset | HDRNA | NDB |  |
| 1 | A(CC) +:WC/H:WT | Type I | 318 | 8 | 9 | rRNA (recurrent) |
| 2 | G(CC) H:+T/W:WC | Type II | 155 | 5 | 9 | rRNA (recurrent), synthetic RNA construct |
| 3 | C(AA) W:+C/S:SC | Type II | 174 | 6 | 6 | rRNA (recurrent) |
| 4 | G(Aψ) H:+T/W:WC<sup>a</sup> | Type II | 19 | 5 | 7 | tRNA |
| 5 | G(CC) H:+C/W:WC | Type II | 5 | 4 | 5 | synthetic RNA construct |
| 6 | G(AC) H:zC/W:WC | Type II | 106 | 3 | 1 | rRNA (non-recurrent) |
| 7 | A(GA) +:HC/S:SC | Type I | 28 | 1 | 3 | rRNA (recurrent) |
| 8 | G(AC) H:+T/W:WC | Type II | 6 | 1 | 3 | tRNA |
| 9 | G(CC) z:WC/W:WC | Type I | 5 | 0 | 4 | synthetic RNA construct |
| 10 | A(CA) +:WC/S:SC | Type I | 16 | 0 | 3 | rRNA (non-recurrent) |
| 11 | C(CG) W:+C/W:WC | Type II | 7 | 0 | 3 | rRNA (non-recurrent), synthetic RNA construct |
| 12 | A(GU) +:HC/S:SC | Type I | 1 | 1 | 1 | synthetic RNA construct |
| 13 | G(CA) W:+C/S:WT | Type II | 19 | 0 | 1 | rRNA (non-recurrent) |
| 14 | G(CA) W:+C/S:ST | Type II | 14 | 1 | 0 | rRNA (non-recurrent) |
| 15 | C(GG) +:WC/W:WC | Type I | 9 | 0 | 1 | rRNA (non-recurrent) |
| 16 | C(AC) W:+C/S:HC | Type II | 2 | 1 | 0 | rRNA (non-recurrent) |
| 17 | C(CG) +:WT/H:HT | Type I | 1 | 0 | 1 | synthetic RNA construct |
| 18 | A(CU) +:WC/H:WC | Type I | 0 | 0 | 1 | rRNA (non-recurrent) |
| 19 | C(GG) +:HC/W:ST | Type I | 0 | 0 | 1 | rRNA (non-recurrent) |
| 20 | G(GU) z:HT/W:WC | Type I | 6 | 0 | 0 | rRNA (non-recurrent) |

<sup>a</sup> $\psi$ stands for the modified nucleobase pseudouracil.

### Both Type I and Type II Protonated Triples Occur Recurrently in Ribosomal RNAs

The most frequently occurring protonated base triple is A(CC) +:W Cis/H:W Trans (*System 1*). Interestingly, it is a Type I protonated triple. Out of all the protonated triples found in the larger dataset, ∼33% belongs to *System 1*. Analysis of its occurrence contexts reveals that this particular triple occurs as a part of many conserved motifs in different RNAs (Figure 2 and Table S6). It occurs as a part of a conserved 5 way junction loop in the Domain V of the 23S rRNA of prokaryotes, archea and eukaryotes (25S/28S rRNA) (Figure 2B, 2C). In the 23S rRNA of archea, it is also found in an internal loop of Domain II (Figure 2D, 2E). In addition to that, in the mitochondrial 16S rRNA of mammals this particular protonated base triple occur at a 3 way junction loop (Figure 2F). These observations therefore rationalize the high occurrence frequency of *System 1* in the RNA crystal structure data sets.

**Figure 2:**
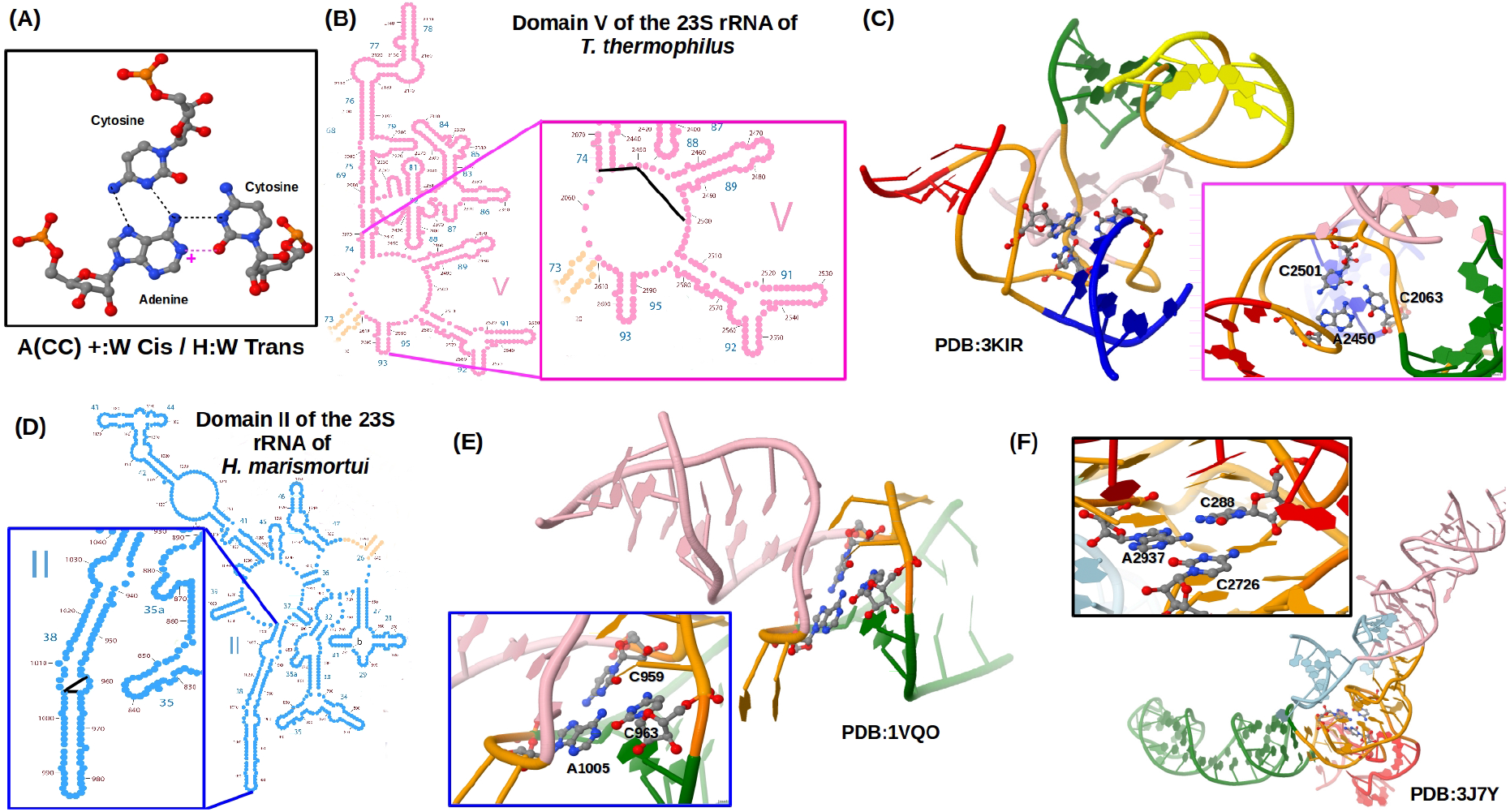
Context of occurrences of A(CC) +:W Cis/H:W Trans triple (*System 1*). (A) Geometry of the base triple. Inter-base hydrogen bonds are shown in broken lines (magenta color represents the proton mediated hydrogen bond). (B) 2D representation of the Domain V of the 23S rRNA of *T. thermophilus*. Enlarged view of the junction loop containing *System 1* is shown with the black lines representing the base-base connections. (C) 3D structure of the junction loop (shown in orange color) connecting 5 helices (shown in green, yellow, blue, red and light-pink color). (D) 2D representation of the Domain II of the 23S rRNA of *H. marismortui*. Enlarged view of the internal loop containing *System 1* is shown with the black lines representing the base-base connections. (E) 3D structure of the internal loop (shown in orange color) between the helical regions (shown in green and light-pink color). (F) 3D structure of the junction loop in mitochondrial 16S rRNA of *H. sapiens* that contains the above mentioned base triple.

The second most frequent protonated base triple G(CC) H:+ Trans/W:W Cis (*System 2*) is also observed in different functional RNAs. For example, it is found (a) in the 3 way junction loop of the 5S rRNA of *T. thermophilus* and *H. morismortui*, (b) in an internal loop in Domain III of the 23S rRNA of *S. cerevisiae*, (c) in the junction loop connecting the P1.1 and P4 helices in HDV ribozyme and (d) in different viral RNA psuedoknots. These occurrence contexts are illustrated in Figure S1.

Unlike the top two most frequent systems, the C(AA) W:+ Cis/S:S Cis triple (*System 3*) does not occur in internal or junction loops, rather it mediates loop-loop interactions. In some prokaryotic 23S rRNAs (*T. thermophilus* and *D. radiodurans*) it connects an internal loop of Domain II with a hairpin loop of Domain V. Again in some eukaryotic 18S rRNAs (*S. cerevisiae* and *P. falciparum*) it mediates the interaction between one bulge loop and one GNRA tetra loop (in this case GAAA tetra loop) in Domain C. A similar occurrence context for *System 3* is also observed in the mitochondrial 16S rRNA of some mammals (*S. scorfa* and *H. sapiens*). These occurrence contexts are illustrated in Figure S2.

In addition to the top three most frequently occurring systems, the A(GA) +:HC/S:SC triple (*System 7*) also occur recurrently in ribosomal RNAs. Like *System 3*, they also bring together distant loops. For example, they mediate loop-loop interaction between one internal loop in Domain 5′ with another internal loop in Domain 3′m of the 18S rRNA of different species (Table S6). Note that out of these four systems that occur recurrently in rRNAs, *System 1* and *System 7* are Type I protonated triples, whereas *System 2* and *System 3* are Type II protonated triples. Other protonated triples are also detected in different ribosomal RNAs, where they take part in the formation of internal loops (*System 13*, *System 14*, *System 16*), bulge loops (*System 6*), multi junction loops (*System 15*, *System 19*, *System 20*), double helical regions (*System 11*, *System 18*) and loop-loop interactions (*System 10*). However, unlike System 1, 2, 3 and 7, these occurrences are not recurrent in the datasets (Table S7 and Figure S3). The fact that protonated triples can occur in various structural environments, as evident from the above analysis, underline their multifaceted contributions in determining the structural landscape of RNA.

### In tRNAs Protonation and Post-transcriptional Modifications Occur Interchangeably in A Conserved Triple

Two of the Type II protonated triples are found in tRNAs where they mediate the interaction between D-arm and variable loop. They are *System 4* (G(AU) H:+T/W:WC) and *System 8* (G(AC) H:+T/W:WC). *System 4* is composed of U13, G22 and A46 residues. The base triple composed of these three residues is one of the frequently observed higher order interaction in tRNA.^107^ Both the residues 13 and 46 are also prone to post-transcriptional modifications. ^36,108^ Therefore, we observe at least four variations of the 13-22-46 triple in known tRNA crystal structures. The generally observed variation is *Variant 1*, where it is composed of cytosine and neutral guanine at residue 13 and 46, respectively (Figure 3A). In *Variant 2* these two residues undergo covariation, i.e. when C13 gets substituted by U13, the G46 gets concomitantly substituted by A46 and *vice-versa*. To maintain the overall geometry of the triple, the A46 gets protonated at its Watson-Crick edge (Figure 3B). In *Variant 3* (Figure 3C), the residue G46 gets post-transcriptionally modified to 7-methylated guanine (m^7^G). Note that m^7^G is also positively charged. Whereas in *Variant 4* (Figure 3D), the triple has residue C13 modified to pseudouracil (*ψ*) and subsequently residue G46 gets replaced by protonated adenine (similar to *Variant 2*).

**Figure 3:**
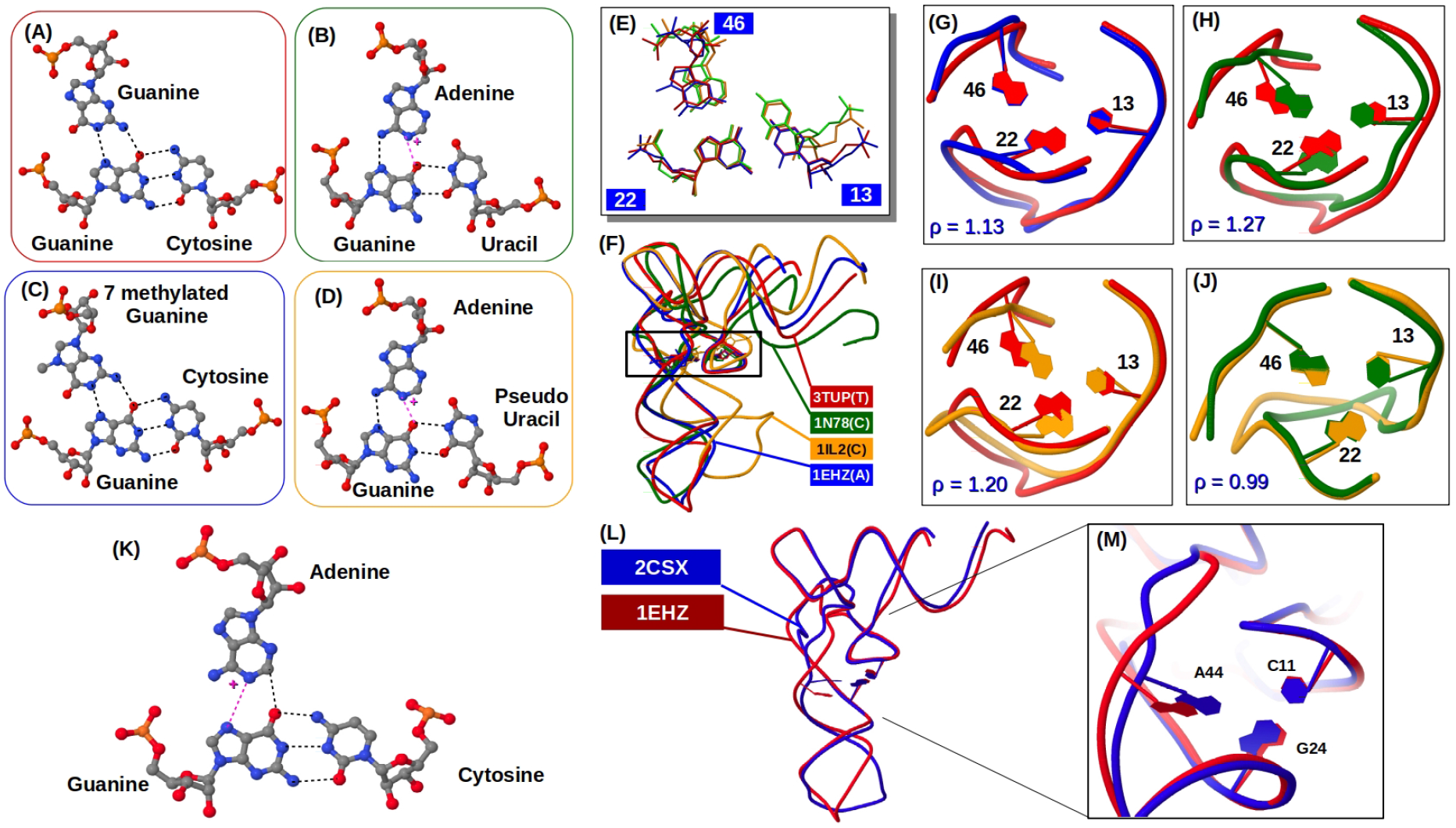
Context of occurrences of the important protonated triples found in tRNAs. **System 4 (G(AU) H:+T/W:WC) :** *System 4* is a tertiary interaction between residues 13, 22 and 46 of tRNA. (A-D) Geometries of four different variants of the base triples. Inter-base hydrogen bonds are shown in broken lines (magenta color represents the proton mediated hydrogen bond). (E) Structural alignment of the four different base triples with respect to the central base 22G. Color scheme is the same labeled in the panel(F). (F) Structural alignment of four different tRNA chains with four different combination of 13-22-46 interaction; 3TUP (13C-22G-46G), 1EHZ (13C-22G-46m^7^G), 1N78 (13U-22G-46A) and 1IL2 (13*φ*-22G-46A). (G-J) Structural alignment of neighborhood of the 13-22-46 triple. ρ represents the RMSD between the aligned structural fragments. **System 8 (G(AC) H:+T/W:WC):**The triple corresponding to *System 8* is present in the pdb 2CSX, but is absent in the pdb 1EHZ. (K) Geometry of the base triple. (L) Structural alignment of the complete tRNA chains of 1EHZ and 2CSX. (M) Enlarged vies of the region around the 11-24-44 triple. Difference in the backbone fold near residue 44 is visible.

Figure 3E shows structural superposition of these four variants of the base triple. The noticeable difference is due to the fact that, in *Variant 2* (green) and *Variant 4* (yellow), the central guanine (G22) forms a wobble G:U pair with the U13/*ψ*13, whereas in *Variant 1* (red) and *Variant 3* (blue) it forms a canonical G:C pair. Figure 3F shows the structural alignment of the complete tRNA molecules containing these four variants of the 13-22-46 triple. A closer view is obtained by performing structural alignment of only the neighborhood of the 13-22-46 triple (Figure 3G-3J). As illustrated in Figure 3G, modification of G46 to m^7^G46 causes noticeable difference in the backbone folding near the residue 46 (RMSD (ρ) = 1.13 Å). The difference further increases on substitution of G46 to A46 (ρ = 1.20 Å in Figure 3I and ρ = 1.27 Å in Figure 3H). Earlier it was shown that guanine N7 protonation^53^ and the archaeosine modification^109^ play similar roles in determining the stability of the Levitt base pair,^110^ which is a conserved interaction between G15 and C48 residues in tRNA. Here also we observe that both nucleobase protonation and post-transcriptional modification at residue 46 causes noticeable change in the local backbone orientation. Though the exact nature of the changes are not identical, it may be argued that nucleobase protonation can compete with post-transcriptional modifications in modulating the folded structure of RNA. It is also noteworthy that modification of U13 to *ψ*13 have a significant influence on the shape of the local backbone (Figure 3J).

The triple corresponding to *System 8* (C11-G24-A44, Figure 3K) is not conserved in tRNA. Structural alignment of a tRNA chain containing the C11-G24-A44 triple with the one lacking it, reveals that formation of the C11-G24-A44 triple results in significant change in the folding of the tRNA chain near the residue 44 (Figure 3L, 3M). These observations highlight the significance of positively charged nucleobases (caused by post-transcriptional modification or direct protonation) on the overall folding of RNA molecules.

### Protonated Triples Stabilize Tertiary Structures of Synthetic RNA Constructs

Occurrences of protonated base triples are not limited to natural RNA molecules only. Six out of the twenty systems are also found in synthetic RNA constructs. They participate in the formation of different tertiary motifs (Table S8), e.g. pseudoknot (*System 2*), junction loops (*System 12*) and helix-helix interactions (*System 17*). In a synthetic tRNA CCA acceptor (PDB Id: 3VJR) and a synthetic vitamin B12 binding RNA aptamer (PDB Id: 1ET4), protonated triples are found to construct tertiary contacts between two different chains (Figure S4). These examples underline the putative nano-biotechnological applications of these protonated triples.

### Both Type I and Type II Protonated Base Triples Are Intrinsically Stable

Occurrence of protonated triples in diverse secondary structural motifs in RNA drives us to probe their energetic contribution in stabilizing those motifs which, in turn, compensate for the energetic cost of protonation. So we calculated the intrinsic geometries and interaction energies of the isolated triples using QM based methods. Figure 4 and Figure S5 illustrate the deviation of the M05-2X/6-31++G(2d,2p) level optimized geometries from their respective crystal geometries. Note that one of the bases has been considered as protonated in all these systems. On geometry optimization, the 4 systems illustrated in Figure S5 diverge significantly from their respective crystal geometries (RMSD(1) > 1.8 Å and RMSD(2) > 2.0 Å). Rest 16 systems (7 Type I and 9 Type II) converge to geometries which are relatively close to their respective native geometries (Figure 4). Therefore, for majority of the cases, the protonation hypothesis can explain the stability of these triples within the crystal environment. It may however be noted that, although the geometry of interaction between the three bases remains unchanged, the native inter-base hydrogen bonding pattern may alter in some systems. Two such examples are illustrated in Figure S6.

**Figure 4:**
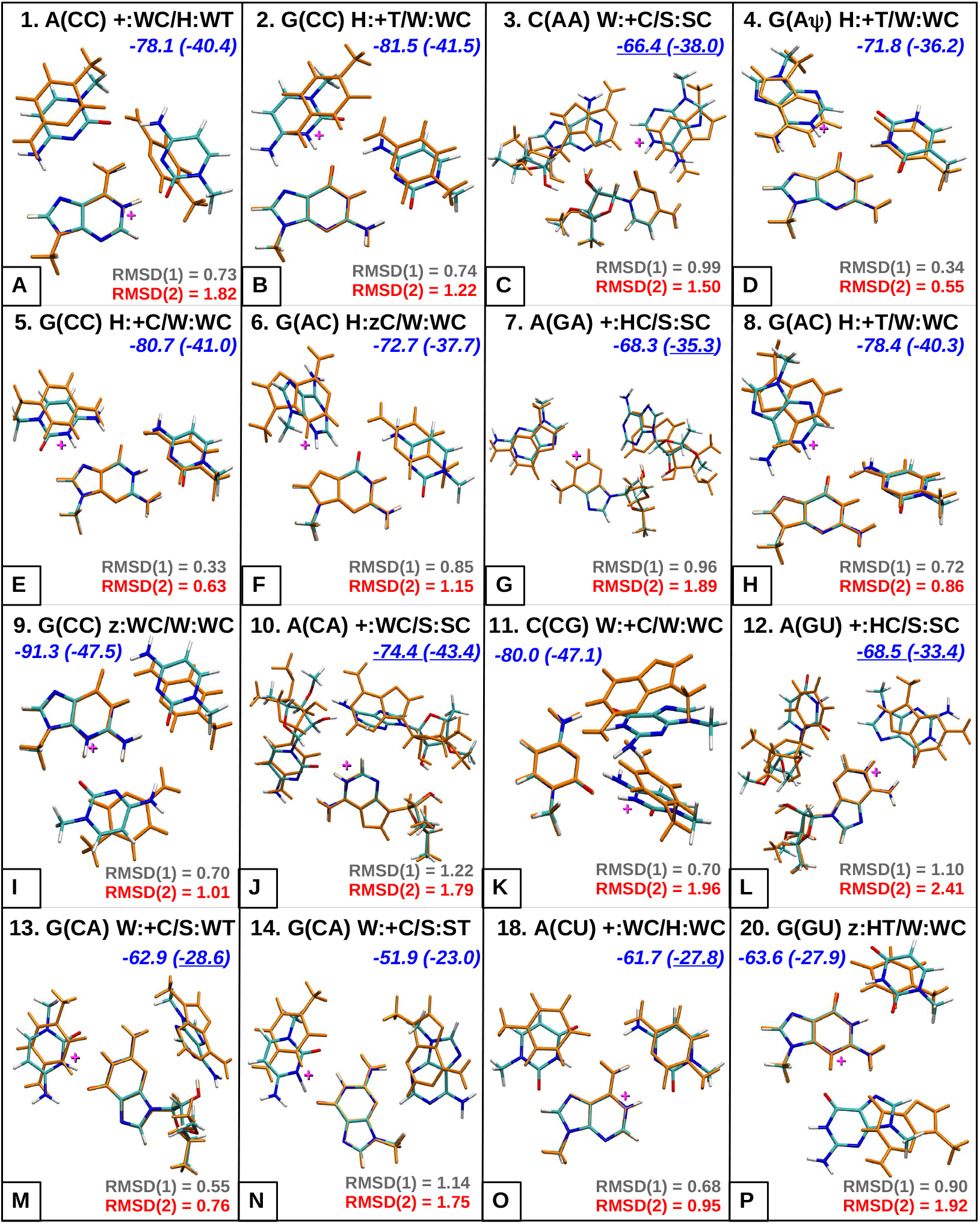
Protonated triples which do not diverge significantly from their respective crystal geometries on geometry optimization performed at M05-2X/6-31++G(2d,2p) level. Superposition of the final optimized geometry (represented in CPK color scheme) over the initial crystal geometry (represented in orange color), with respect to the central base is shown. MP2/aug-cc-pVDZ level interaction energies of the optimized triples are reported (blue text) in kcal mol^−1^. The values reported within parenthesis indicate the solvent phase interaction energies. The MP2 level energies that are underlined are calculated using 6-31+G(d,p) basis set to avoid convergence error. RMSD(1) and RMSD(2) values are reported in Å in gray and red colored text, respectively.

The 4 nonconforming systems (Systems 15, 16, 17 & 19) shown in Figure S5 has two bifurcated triples (*System 15* and *System 19*). In bifurcated triples two terminal bases interact with the same edge of the central base. Interestingly another bifurcated triple studied here (*System 11*) is stable under geometry optimization (Figure 4K) at M05-2X/6-31++G(2d,2p) level. Analysis of the context of occurrence of these bifurcated base triples reveal that both *System 15* and *System 19*, the unstable bifurcated triples, are found in multijunction loops in eukaryotic 28S rRNA (Table S7). Whereas, *System 11*, the stable bifurcated triple, occur within double helical stretches in 28S rRNA of *S scorfa*, Signal Recognition Particle (SRP) RNA of *P. horikoshii* and synthetic double stranded RNA fragments (e.g. 2GJW). Similarly, as mentioned above, another unstable system viz. *System 17* is involved in mediating helix-helix interactions in synthetic RNAs.

Therefore, we argue that geometries of protonated triples that build tertiary interactions between distant motifs are largely constrained by their respective crystal environments. In the absence of those constraints during geometry optimization performed in isolation, they diverge from their respective crystal geometries. Such environmental constraints are rare in protonated triples that do not build tertiary interactions between distant motifs but contribute in shaping the secondary structural motifs.

### Weaker Triples May Occur Recurrently Due to Their Specific Functional Requirements

Interaction energies (calculated at MP2/aug-cc-pVDZ level) of the protonated base triples reported in Figure 4 and their occurrence frequencies reported in Table 1. They collectively suggest that the frequency of occurrence and intrinsic stability of these base triples are not correlated. For example, the rarely occurring (5 instances in the larger database) *System 9* has the highest interaction energy (−91.3 kcal mol^−1^ in gas phase), whereas the frequently occurring (174 instances in the larger database) *System 3* is relatively weak (−66.4 kcal mol^−1^ in gas phase) and comes at the 12^*th*^ position in the order of interaction energy.

We also observe that out of the 16 stable protonated triples (Figure 4) there are 5 triples (*System 3*, *7*, *10*, *12* and *14*) which contain a SS base pair (*cis* or *trans*). All of them have higher RMSD values and lower interaction energies compared to other stable protonated triples. This implies that they are intrinsically flexible. This extra flexibility of these triples is consistent with earlier QM based analysis of RNA base pairs which have suggested that SS base pairs are relatively more flexible than base pairs of other geometries.^20,111^ Also note that except *System 14*, the other four systems involve interbase hydrogen bonding through the dynamic O2′ atom of the sugar moiety. Interestingly, despite being relatively weak, systems 3, 7 and 10 occur recurrently in rRNAs. In rRNAs they participate in bringing together distant loops (Table S6 and S7), whereas, other protonated triples mostly get involved in formation of internal loops and multi-junction loops. *System 12* is also observed in multi-junction loops in synthetic RNAs. So adding up to our earlier argument, we further propose that the intrinsic flexibility of these base triples with SS pairs are utilized in RNA to construct tertiary contacts between distant motifs. Their functional requirement justifies their recurrent occurrences in RNA.

### Protonated Base Triples Are Stabilized by a Wide Variety of Nonbonding Interactions

In Figure 4 we have reported the interaction energies at MP2 level. We also calculated the interaction energies using two different DFT functionals, viz. M05-2X and ωB97XD (Figure 5A and Table S9), as they treat the dispersion interactions in two different ways. M05-2X functional is parametrized for middle range dispersion interactions, whereas corrections for long range dispersion interactions are explicitly incorporated in ωB97XD functional via Grimme’s DFT-D formalism. Note that DFT level interaction energies reported here include both deformation energy and BSSE corrections, whereas the reported MP2 level interaction energies include only deformation energy correction. Results obtained for both the DFT functionals are consistent with each other (Figure 5A). However, interaction energies obtained using ωB97XD are on average 3.4 kcal mol^−1^ higher than those obtained using M05-2X functional. This highlights the importance of long range dispersion forces in these systems. Notably, all the four systems that have S:S Cis base pairs (systems 3, 7, 10 and 12) show higher differences between ωB97XD and M05-2X level interaction energies. For systems 3, 7, 10 and 12 the differences are 6.7, 5.9, 6.7 and 6.2 kcal mol^−1^, respectively. So for the systems containing interacting sugar moieties, long range dispersion forces become even more important.

**Figure 5:**
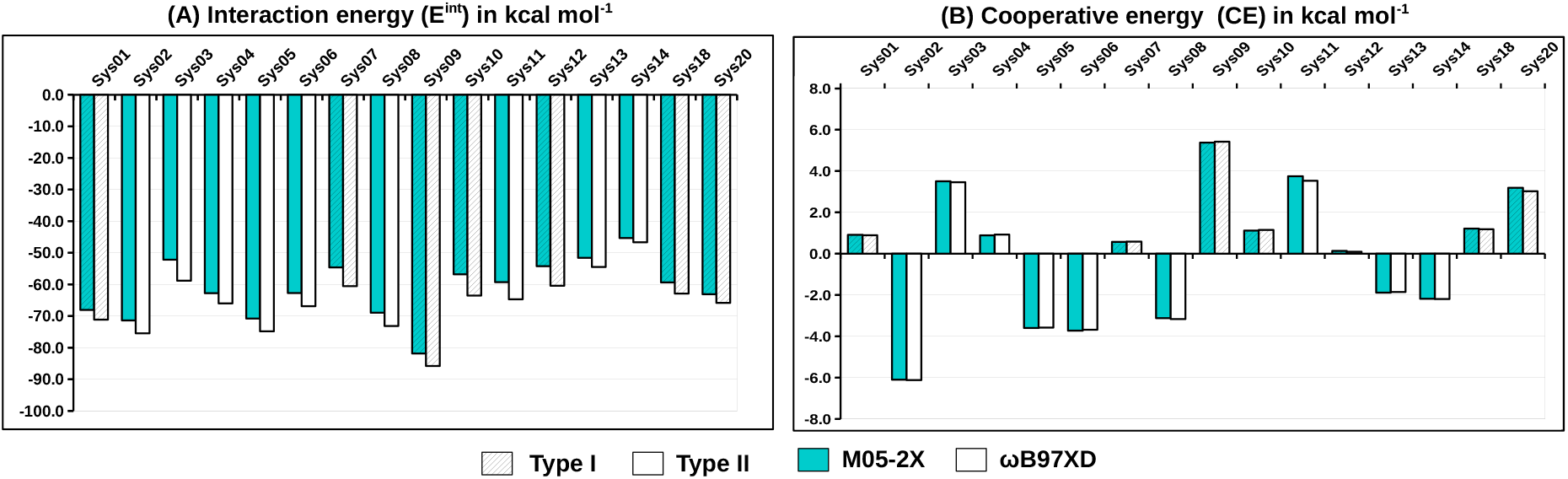
(A) Comparison of the interaction energies and (B) comparison of the cooperative energies of different protonated base triples. Interaction energies and cooperative energies have been calculated using two DFT functionals – (cyan) M05-2X and (white) ωB97XD with 6-311++G(2df,2pd) basis set. Bars corresponding to the Type II triples contain oblique stripes.

Nevertheless, the relative order of DFT level interaction energies remain consistent with MP2 level results, except for *System 10* and *System 11*. For these two systems the differences between MP2 and DFT level interaction energies are remarkably high. Difference between MP2 level and M05-2X level interaction energies for *System 10* and *System 11* are 17.6 kcal mol^−1^ (10.9 kcal mol^−1^ at ωB97XD functional) and 20.7 kcal mol^−1^ (15.7 kcal mol^−1^ at ωB97XD functional), respectively. Probing the origin of such significant difference is beyond the scope of this manuscript. However, QTAIM analysis of these two systems reveal that in addition to inter-base hydrogen bonds, they are stabilized by other nonbonding interactions. As illustrated in Figure S7C, the C-H‖ π type interaction between C2-H group of the central adenine and aromatic ring of the other adenine in *System 10* is characterized by one bond path for which the corresponding BCP has ∇^2^ρ > 0. Again for the case of *System 11* the π–π stacking interaction between the aromatic rings of the two terminal bases is characterized by a number of bond paths for which the corresponding BCPs have ∇^2^ρ > 0 (Figure S7D). So a careful benchmarking of DFT functionals is needed to study these systems stabilized by different types of nonbonding interactions.

It was argued earlier that charge-dipole interaction is an important factor in the overall stability of a protonated base pairs.^42^ Here we observe that the triples composed of high dipole moment bases (guanine and cytosine) have higher interaction energies compared to the systems that are composed of low dipole moment bases (adenine and uracil). For example, gas phase interaction energies of the systems that are composed of one guanine and two cytosine bases (systems 2, 5, 9 and 11) are greater than 80 kcal mol^−1^ (Figure 4) at MP2/aug-cc-pVDZ level. It is non-trivial to determine exact energetic contribution of charge-dipole interactions. However, we can qualitatively argue that protonated triples, which have the scope for higher charge-dipole interactions, are likely to have higher interaction energies.

### Formation of Type II Triples Are Energetically Cooperative, Whereas Type I Triples Are Energetically Anti-cooperative

It is interesting to note that except *System 1*, which has recurrent occurrences in different conserved multi junction loops in different rRNAs, other Type I protonated triples (systems 7, 9, 10, 12, 18 and 20) have remarkably low occurrences in RNA crystal structure datasets (Table 1), despite their high interaction energies (Figure 4). To understand on what basis Type II triples are discriminated in RNA we analyzed their cooperative energies (CEs). For a base triple, the corresponding CE is the difference between its interaction energy and sum of the interaction energies of its three constituent base-base interactions. All the Type I triples have CE > 0 and all the Type II triples have CE < 0 (except systems 3 and 4), as shown in Figure 5. The interaction energies calculated using M05-2X and ωB97Xd functional with 6-311++G(2df,2pd) basis set, and their corresponding cooperative energies (CE), are illustrated in Figure 5. Although out of the two functionals, interaction energies calculated using ωB97XD functional are larger for all the cases, we found that the cooperative energies remain independent of the choice of functional. Similar trend was also reported earlier.^103^

From Figure 5 and Table S9 we found that formation of protonated triples of Type I are energetically anti-cooperative. That means, formation of the *Class I* type protonated base pair, between the central base and the first terminal base, does not favor the formation of the second base pair between the central base and the second terminal base, and *vice versa*. The first possibility has been examined by analyzing the charge redistribution at the free edges due to formation of the *Class I* type protonated base pair (Table S10). In principle, increase in the electronic charge density over a hydrogen bond (H-bond) acceptor site and decrease in the electronic charge density over a H-bond donor site improve their respective H-bonding potentials. The opposite change will worsen the H-bonding potentials. Figure 6 illustrates the observations taking one example of Type I (*System 9*) and one example of Type II (*System 2*) triples. Note that, out of the 20 systems, *System 2* is the most cooperative (CE = −6.0 kcal mol^−1^) and *System 9* is the most anti-cooperative (CE = 5.4 kcal mol^−1^) triple (Figure 5). In both the cases, the central base guanine forms a protonated base pair with one of the terminal cytosines. The protonated base pair involves the sugar edge of guanine in *System 9* (G:C z:W Cis) and involves the Hoogsteen edge of guanine in *System 2* (G:C H:+ Cis), respectively. So in both the cases, the Watson-Crick edge of guanine interacts with the second cytosine. The change in the electron density over the three H-bonding sites of the Watson-Crick edge caused by the formation of *Class I* protonated base pair is characterized by the term %Δq, which is defined as,

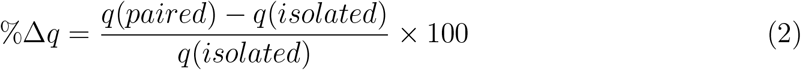

where, q(isolated) and q(paired) indicate the partial charge obtained from Natural Population Analysis at a site before and after the base pairing takes place, respectively. The protonated base pair formation weakens the H-bonding potential of O6 of guanine in *System 9* (Figure 6E). Whereas, in *System 2* (Figure 6F) H-bonding potential improves at all the three sites. The same trend is observed for other systems (Table S10). That is due to the formation of the protonated base pair, the central bases belonging to Type I triples undergo weakening of the H-bonding potential at one or more free sites. On the other hand, the central bases belonging to Type II triples undergo strengthening of the H-bonding potential at all the free sites. This further leads to formation of anti-cooperative triples for Type I and cooperative triples for Type II. Note that this analysis is not possible for System 11 (C(CG) W:+C/W:WC), since it is a bifurcated triple.

**Figure 6:**
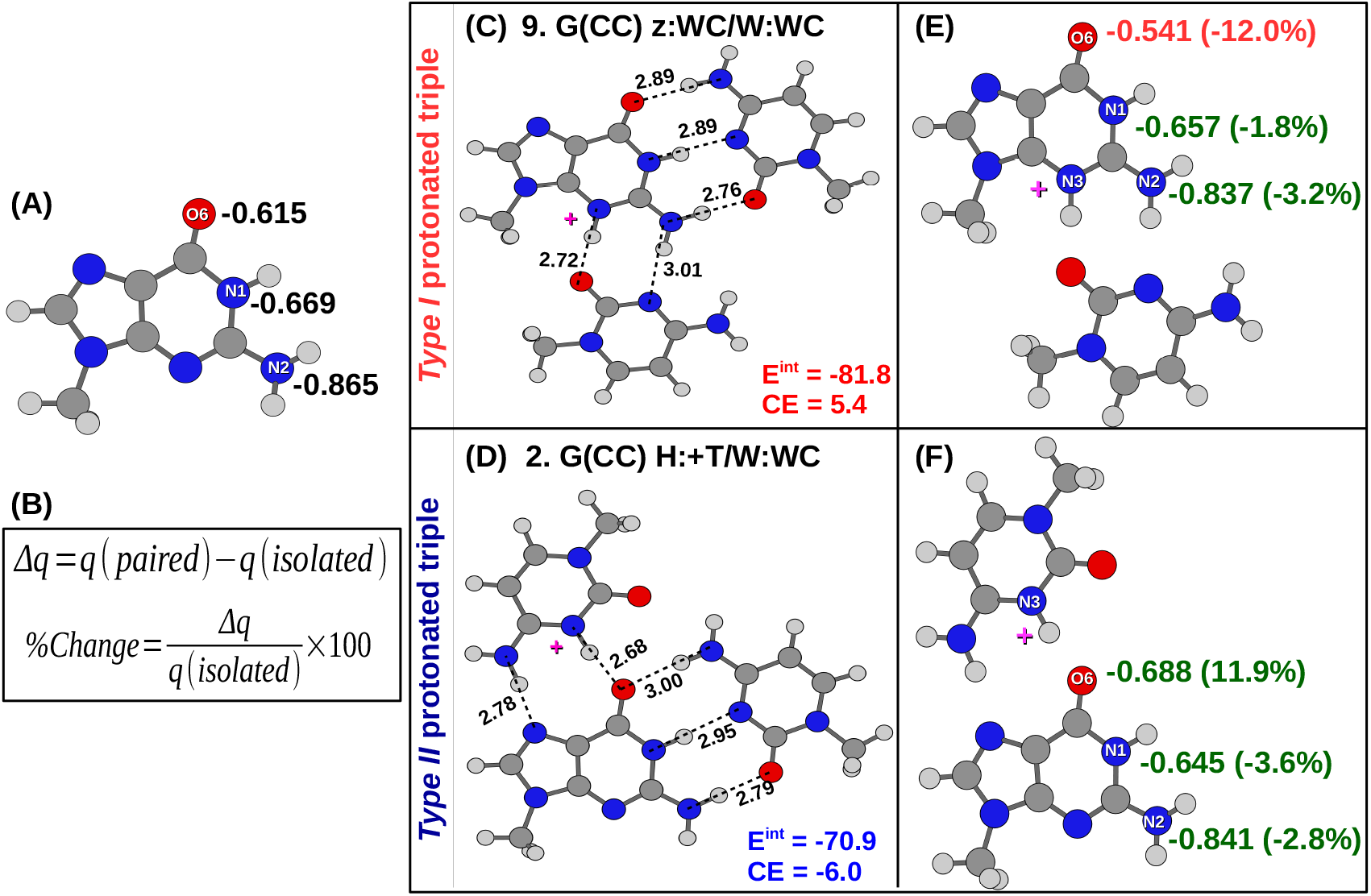
(A) Partial charges at the Watson-Crick edge of the isolated guanine. (B) Definition of %Δq. (C) Optimized geometry of *System 9*. (D) Optimized geometry of *System 2*. Partial charges at the Watson-Crick edge of guanine of the (E) G:C z:W Cis pair and (F) C:G +:H Cis pair. Partial charges are reported in a.u. (fraction of the electronic charge). Hydrogen bonds are represented as broken lines. Hydrogen bond donor-acceptor distances are reported in Å. Interaction energies (E^*int*^) and cooperative energies (CE) are reported in kcal mol^−1^. All the calculations are performed using using M05-2X functional.

There are two exceptions in this general trend. *Systems 3* (Figure S2D) and *System 4* (Figure 3D) are Type II triples but they are anti-cooperative. However, the proton mediated base pairing improves the H-bonding potential at all the free sites in these two systems. We note that, unlike other Type II systems, in these two systems the third base not only interacts with the central base, it also interact with the protonated terminal base through a C-H… O/N type H-bond. This is illustrated in Figure S7A and S7B. However, understanding the rationale behind such anti-cooperative binding requires further investigation and is beyond the scope of this work.

To summarize, we have identified 19 different protonated base triples occurring in two nonredundant sets of RNA crystal structure. Another triple considered in this work occurs frequently in a larger collection of RNA structures. 10 of them are Type I triples that have the protonated base at the central position. Other 10 triples are of Type II which have the protonated base at one of the terminal positions. We observe that, both Type I and Type II triples can occur recurrently in conserved structural motifs in rRNAs. Out of the recurrently occurring base triples some are involved in building tertiary contacts between distant motifs. Others are involved in shaping up the secondary structures. In the former case the triples involved are composed of SS pairs. We found that these triples that bring together distant motifs deviate largely from their respective crystal structures on gas phase geometry optimization. This implies that they remain structurally constrained within the crystal environment. Also their interaction energies are relatively low. Therefore we argue that intrinsic flexibility of these base triples with SS pairs are tailor made for supporting tertiary interactions between distant structural motifs in RNA. Such specific functional requirements justify their recurrent occurrences. We further argue that nucleobase protonation can compete with post-transcriptional modifications to modulate the folding of RNA since we find that nucleobase protonation and post-transcriptional modifications occur interchangeably at a conserved triple in tRNA. Analyzing the interaction energies of protonated triples obtained using two different DFT functionals, and also using MP2 theory, we conclude that, in addition to inter-base hydrogen bonds, multiple nonbonding interactions, such as, charge-dipole interactions, long range dispersion interactions, etc play important roles in stabilization of different protonated triples. Interestingly, formation of Type I triples are energetically anti-cooperative, whereas Type II triples are cooperative. In Type II triples charge redistribution on the free edges of the central base, caused by formation of the *Class I* protonated base pair, favors binding of the third base. This is the opposite for Type I triples. Consequently we observe that Type II triples occur more frequently in RNA crystal structure datasets.

## Conclusion

In general it is argued that the energy released, due to participation of protonated nucleobases in different inter-nucleotide interactions, compensates for the energetic penalty for nucleobase protonation at physiological pH. However the detailed accounting remains elusive for the conventional biophysical methods.^34^ Our investigations using computational methods shed some new light on this long standing problem.

The simplest form of such inter-nucleotide interactions, which has also been explored widely, is the *Class I* type base pairing interaction between a protonated base and a neutral base. In this work we have shown that the *Class I* protonated base pairs are involved in at least 58 different types of base triples in RNA. In one category of these triples the protonated base of the *Class I* protonated base pair interacts with the third base (Type I), and in the other category the neutral base of the protonated pair interact with the third base (Type II). Out of these two categories of protonated base triples, formations of only the later are energetically cooperative and hence are favored in RNA.

We have also argued that geometry and structural flexibility of some protonated triples are tailor made for specific conserved RNA motifs. Therefore they can also frequently occur in RNA, despite being weak or energetically anti-cooperative. Unraveling of the fact that most of such weak and flexible protonated triples connect distant structural motifs in RNA is expected to aid de-novo design and prediction of RNA 3D structures.^112^ With the growth in the utilization of stable protonated base pairs in RNA nanotechnology^113^ our results, showing existence of different protonated base triples with varied energetic stability, should be useful for scientists working on design of novel RNA based nano-devices like pH biosensors.

Chemically modified nucleosides in RNA have recently emerged as critical post-transcriptional regulators of the gene expression process. ^114,115^ It is even found that certain RNA modification pathways are misregulated in human cancers and could be plausible targets of cancer therapy.^116^ From our analyses it is evident that like post-transcriptional modifications, nucleobase protonation in RNA span all three phylogenetic domains (Archaea, Bacteria, and Eukarya) and are ubiquitous across RNA types, including mRNA (riboswitches), tRNA, rRNA, etc. We also argue that nucleobase protonation may occur interchangeably with post-transcriptional modifications and have similar consequences on RNA’s structure. However, the possibility of involvement of protonated nucleobases in gene regulation has not been explored thoroughly. Note that, post-transcriptional modifications are permanent changes to the nucleotides, whereas nucleobase protonation can be considered as a reversible response to environmental or cellular stimuli leading to conformational switching in RNA. Interestingly, Baisden and co-workers^43^ have recently proved that maturation of oncogenic microRNA-21 strongly depends on such pH driven conformational switching events in its precursor. Using NMR relaxation dispersion spectroscopy and mutagenesis they have discovered a transiently occuring but kinetically stable excited state of the precursor microRNA, where formation of a protonated A(+):C W:W Cis pair reshuffles the entire secondary structure including the Dicer cleavage site. This opens up avenues for exploring direct and indirect contributions of nucleobase protonation in regulation of gene expression. Our results revealing recurrent and functionally driven occurrences of protonated base triples in different RNA motifs and justifying the source of their thermodynamic stability provide useful insights required for scouting those avenues.

## Acknowledgement

AH acknowledges CSIR for SRF support. A.M. and A.H. thank DBT, Government of India, project BT/PR-14715/PBD/16/903/2010 for partial funding and financial support. A.M. and D.B. thank DBT, Government of India, project BT/PR-11429/BID/07/271/2008 for supporting computational infrastructure. Authors also thank (a) Dr. Soumen Saha, Nagoya University, Nagoya, Japan for sharing his experience in carrying out CE calculations and (b) Dr. Sohini Bhattacharya, Dr.Reddy’s Institute of Life Sciences, Hyderabad, India for helping out in bioinformatic analysis.

## Supporting Information Available

Base pairing nomenclature used in the manuscript, Count of total number of possible base pairs, PDB Ids of the RNA structures belonging to HD-RNAS, NDB and the ‘larger-dataset’, Table S1-S9 and Figure S1-S6. Table S1: occurrences of protonated triples in the ‘larger dataset’, Table S2: occurrences in NDB dataset, Table S3: occurrences in HDRNAS dataset, Table S4: details of protonated triples selected for QM studies, Table S5: benchmarking of DFT functionals, Table S6: details of recurrent occurrences in rRNAs, Table S7: details of non-recurrent occurrences in rRNAs, Table S8: details of occurrences in synthetic RNAs, Table S9: different components of interaction energies, Table S10: partial charge redistribution at the free edges of the neutral base of a *Class I* protonated base pair. Figure S1: context of occurrences of *System 2*, Figure S2: context of occurrences of *System 3*, Figure S3: context of occurrences of *System 6* and *System 14*, Figure S4: context of occurrences of *System 5* and *System 9*, Figure S5: optimized geometries of systems 15, 16, 17 & 19, Figure S6: changes in inter-base hydrogen bonding pattern, Figure S7: results of AIM analysis.

## References

(1) Eddy, S. R. Non–coding RNA genes and the modern RNA world. Nat. Rev. Genet. 2001, 2, 919–929.

(2) Cech, T. R.; Steitz, J. A. The noncoding RNA revolution—trashing old rules to forge new ones. Cell 2014, 157, 77–94.

(3) Crick, F. Central dogma of molecular biology. Nature 1970, 227, 561–563.

(4) Cox, M. M.; Nelson, D. L. Lehninger principles of biochemistry, 5th ed.; WH Freeman, 2008.

(5) Lakhotia, S. C.; Mallick, B.; Roy, J. In RNA-Based Regulation in Human Health and Disease; Pandey, R., Ed.; Translational Epigenetics; Academic Press, 2020; Vol. 19; pp 5–57.

(6) He, L.; Hannon, G. J. MicroRNAs: small RNAs with a big role in gene regulation. Nat. Rev. Genet. 2004, 5, 522–531.

(7) Morris, K. V. RNA and the regulation of gene expression: A hidden layer of complexity; Horizon Scientific Press, 2008.

(8) Matera, A. G.; Terns, R. M.; Terns, M. P. Non-coding RNAs: lessons from the small nuclear and small nucleolar RNAs. Nat. Rev. Mol. Cell Biol. 2007, 8, 209–220.

(9) Kikovska, E.; Svärd, S. G.; Kirsebom, L. A. Eukaryotic RNase P RNA mediates cleavage in the absence of protein. Proc. Natl. Acad. Sci. U.S.A. 2007, 104, 2062–2067.

(10) Serganov, A.; Patel, D. J. Ribozymes, riboswitches and beyond: regulation of gene expression without proteins. Nat. Rev. Genet. 2007, 8, 776–790.

(11) Šponer, J.; Bussi, G.; Krepl, M.; Banáš, P.; Bottaro, S.; Cunha, R. A.; Gil-Ley, A.; Pinamonti, G.; Poblete, S.; Jurečka, P.; Walter, N. G.; Otyepka, M. RNA structural dynamics as captured by molecular simulations: A comprehensive overview. Chem. Rev. 2018, 118, 4177–4338.

(12) Woodson, S. A. Compact but disordered states of RNA. Nat. Struct. Biol. 2000, 7, 349–352.

(13) Thirumalai, D.; Lee, N.; Woodson, S. A.; Klimov, D. Early events in RNA folding. Annu. Rev. Phys. Chem. 2001, 52, 751–762.

(14) Woodson, S. A. Metal ions and RNA folding: a highly charged topic with a dynamic future. Curr. Opin. Chem. Biol. 2005, 9, 104–109.

(15) Herschlag, D.; Bonilla, S.; Bisaria, N. The story of RNA folding, as told in epochs. Cold Spring Harb. Perspect. Biol. 2018, 10.

(16) Egli, M.; Saenger, W. Principles of Nucleic Acid Structure; Springer Advanced Texts in Chemistry; Springer-Verlag, 1984; pp P001–556.

(17) Westhof, E.; Fritsch, V. RNA folding: Beyond Watson–Crick pairs. Structure 2000, 8, R55–R65.

(18) Leontis, N. B.; Westhof, E. Geometric nomenclature and classification of RNA base pairs. RNA 2001, 7, 499–512.

(19) Bhattacharya, S.; Mittal, S.; Panigrahi, S.; Sharma, P.; Preethi, S.; Paul, R.; Halder, S.; Halder, A.; Bhattacharyya, D.; Mitra, A. RNABP COGEST: a resource for investigating functional RNAs. Database 2015, 2015, bav011.

(20) Šponer, J. E.; Leszczynski, J.; Sychrovský, V.; Šponer, J. Sugar edge/sugar edge base pairs in RNA: Stabilities and structures from quantum chemical calculations. J. Phys. Chem. B 2005, 109, 18680–18689.

(21) Šponer, J. E.; Špačková, N.; Kulhánek, P.; Leszczynski, J.; Šponer, J. Non-Watson-Crick base pairing in RNA. Quantum chemical analysis of the cis Watson-Crick/sugar edge base pair family. J. Phys. Chem. A 2005, 109, 2292–2301.

(22) Šponer, J. E.; Špačková, N.; Leszczynski, J.; Šponer, J. Principles of RNA base pairing: structures and energies of the trans Watson-Crick/sugar edge base pairs. J. Phys. Chem. B 2005, 109, 11399–11410.

(23) Sharma, P.; Mitra, A.; Sharma, S.; Singh, H.; Bhattacharyya, D. Quantum chemical studies of structures and binding in noncanonical RNA base pairs: The trans Watson-Crick:Watson-Crick family. J. Biomol. Struct. Dyn. 2008, 25, 709–732.

(24) Mládek, A.; Sharma, P.; Mitra, A.; Bhattacharyya, D.; Šponer, J.; Šponer, J. E. Trans Hoogsteen/sugar edge base pairing in RNA. Structures, energies, and stabilities from quantum chemical calculations. J. Phys. Chem. B 2009, 113, 1743–1755.

(25) Sharma, P.; Chawla, M.; Sharma, S.; Mitra, A. On the role of Hoogsteen:Hoogsteen interactions in RNA: Ab initio investigations of structures and energies. RNA 2010, 16, 942–957.

(26) Sharma, P.; Sponer, J. E.; Sponer, J.; Sharma, S.; Bhattacharyya, D.; Mitra, A. On the role of the cis Hoogsteen:Sugar-edge family of base pairs in platforms and triplets — Quantum chemical insights into RNA structural Biology. J. Phys. Chem. B 2010, 114, 3307–3320.

(27) Correll, C. C.; Swinger, K. Common and distinctive features of GNRA tetraloops based on a GUAA tetraloop structure at 1.4 Å resolution. RNA 2003, 9, 355–363.

(28) Bottaro, S.; Lindorff-Larsen, K. Mapping the universe of RNA tetraloop folds. Biophys. J. 2017, 113, 257–267.

(29) Heus, H. A.; Pardi, A. Structural features that give rise to the unusual stability of RNA hairpins containing GNRA loops. Science 1991, 253, 191–194.

(30) Halder, S.; Bhattacharyya, D. RNA structure and dynamics: a base pairing perspective. Prog. Biophys. Mol. Biol. 2013, 113, 264–283.

(31) Cantara, W. A.; Crain, P. F.; Rozenski, J.; McCloskey, J. A.; Harris, K. A.; Zhang, X.; Vendeix, F. A. P.; Fabris, D.; Agris, P. F. The RNA modification database, RNAMDB: 2011 update. Nucleic Acids Res. 2011, 39, D195–D201.

(32) Leonarski, F.; D’Ascenzo, L.; Auffinger, P. Mg^2+^ ions: Do they bind to nucleobase nitrogens? Nucleic Acids Res. 2017, 45, 987–1004.

(33) Westhof, E. Isostericity and tautomerism of base pairs in nucleic acids. FEBS Letters 2014, 588, 2464–2469.

(34) Wilcox, J. L.; Ahluwalia, A. K.; Bevilacqua, P. C. Charged nucleobases and their potential for RNA catalysis. Acc. Chem. Res. 2011, 44, 1270–1279.

(35) Chawla, M.; Oliva, R.; Bujnicki, J. M.; Cavallo, L. An atlas of RNA base pairs involving modified nucleobases with optimal geometries and accurate energies. Nucleic Acids Res. 2015, 43, 6714–6729.

(36) Seelam, P. P.; Sharma, P.; Mitra, A. Structural landscape of base pairs containing post-transcriptional modifications in RNA. RNA 2017, 23, 847–859.

(37) Halder, A.; Roy, R.; Bhattacharyya, D.; Mitra, A. Consequences of Mg^2+^ binding on the geometry and stability of RNA base pairs. Phys. Chem. Chem. Phys. 2018, 20, 21934–21948.

(38) Watson, J. D.; Crick, F. H. Genetical implications of the structure of deoxyribonucleic acid. Nature 1953, 171, 964–967.

(39) Topal, M. D.; Fresco, J. R. Complementary base pairing and the origin of substitution mutations. Nature 1976, 263, 285–289.

(40) Singh, V.; Fedeles, B. I.; Essigmann, J. M. Role of tautomerism in RNA biochemistry. RNA 2015, 21, 1–13.

(41) Rozov, A.; Westhof, E.; Yusupov, M.; Yusupova, G. The ribosome prohibits the G•U wobble geometry at the first position of the codon–anticodon helix. Nucleic Acids Res. 2016, 44, 6434–6441.

(42) Chawla, M.; Sharma, P.; Halder, S.; Bhattacharyya, D.; Mitra, A. Protonation of base pairs in RNA: context analysis and quantum chemical investigations of their geometries and stabilities. J. Phys. Chem. B 2011, 115, 1469–1484.

(43) Baisden, J. T.; Boyer, J. A.; Zhao, B.; Hammond, S. M.; Zhang, Q. Visualizing a protonated RNA state that modulates microRNA-21 maturation. Nat. Chem. Biol. 2021, 17, 80–88.

(44) Wolter, A. C.; Weickhmann, A. K.; Nasiri, A. H.; Hantke, K.; Ohlenschläger, O.; Wunderlich, C. H.; Kreutz, C.; Duchardt-Ferner, E.; Wöhnert, J. A stably protonated adenine nucleotide with a highly shifted pK_*a*_ value stabilizes the tertiary structure of a GTP-binding RNA aptamer. Angew. Chem. Int. Ed. 2017, 56, 401–404.

(45) Pechlaner, M.; Donghi, D.; Zelenay, V.; Sigel, R. K. O. Protonation-dependent base flipping at neutral pH in the catalytic triad of a self-splicing bacterial group II intron. Angew. Chem. Int. Ed. 2015, 54, 9687–9690.

(46) Bink, H. H. J.; Hellendoorn, K.; van der Meulen, J.; Pleij, C. W. A. Protonation of non-Watson–Crick base pairs and encapsidation of turnip yellow mosaic virus RNA. Proc. Nat. Acad. Sci. 2002, 99, 13465–13470.

(47) Gehring, K.; Leroy, J.-L.; Gueron, M. A tetrameric DNA structure with protonated cytosine-cytosine base pairs. Nature 1996, 363, 561–565.

(48) Berger, I.; Kang, C.; Fredian, A.; Ratliff, R.; Moyzis, R.; Rich, A. Extension of the four-stranded intercalated cytosine motif by adenine-adenine base pairing in the crystal structure of d(CCCAAT). Nat. Struct. Mol. Biol. 1995, 2, 416–425.

(49) Mirkin, S. M.; Frank-Kamenetski, M. D. H-DNA and Related Structures. Ann. Rev. Biophys. and Biomol. Struct. 1994, 23, 541–576.

(50) Zain, R.; Sun, J.-S. Do natural DNA triple-helical structures occur and function in vivo? Cell and Mol. Life. Sci. 2003, 60, 862–870.

(51) Nakano, S.-i.; Chadalavada, D. M.; Bevilacqua, P. C. General acid-base catalysis in the mechanism of a hepatitis delta virus ribozyme. Science 2000, 287, 1493–1497.

(52) Bevilacqua, P. C.; Brown, T. S.; Chadalavada, D.; Lecomte, J.; Moody, E.; I., N. S. Linkage between proton binding and folding in RNA: Implications for RNA catalysis. Biochem. Soc. Trans. 2005, 33, 466–470.

(53) Halder, A.; Bhattacharya, S.; Datta, A.; Bhattacharyya, D.; Mitra, A. The role of N7 protonation of guanine in determining the structure, stability and function of RNA base pairs. Phys. Chem. Chem. Phys. 2015, 17, 26249–26263.

(54) Halder, A.; Vemuri, S.; Roy, R.; Katuri, J.; Bhattacharyya, D.; Mitra, A. Evidence for hidden involvement of N3-protonated guanine in RNA structure and function. ACS Omega 2019, 4, 699–709.

(55) Gleghorn, M. L.; Zhao, J.; Turner, D. H.; Maquat, L. E. Crystal structure of a poly(rA) staggered zipper at acidic pH: evidence that adenine N1 protonation mediates parallel double helix formation. Nucleic Acids Res. 2016, 44, 8417–8424.

(56) Bevilacqua, P. C.; Brown, T. S.; Nakano, S.-i.; Yajima, R. Catalytic roles for proton transfer and protonation in ribozymes. Biopolymers 2004, 73, 90–109.

(57) Verdolino, V.; Cammi, R.; Munk, B. H.; Schlegel, H. B. Calculation of pK_*a*_ values of nucleobases and the guanine oxidation products guanidinohydantoin and spiroiminodihydantoin using density functional theory and a polarizable continuum model. J. Phys. Chem. B 2008, 112, 16860–16873.

(58) Yang, B.; Wu, R.; Rodgers, M. Base-pairing energies of proton-bound homodimers determined by guided ion beam tandem mass spectrometry: Application to cytosine and 5-substituted cytosines. Anal. Chem. 2013, 85, 11000–11006.

(59) Yang, B.; Moehlig, A. R.; Frieler, C.; Rodgers, M. Base-pairing energies of protonated nucleobase pairs and proton affinities of 1-methylated cytosines: Model systems for the effects of the sugar moiety on the stability of DNA i-motif conformations. J. Phys. Chem. B 2015, 119, 1857–1868.

(60) Halder, A.; Halder, S.; Bhattacharyya, D.; Mitra, A. Feasibility of occurrence of different types of protonated base pairs in RNA: a quantum chemical study. Phys. Chem. Chem. Phys. 2014, 16, 18383–18396.

(61) Alkorta, I.; Mata, I.; Molins, E.; Espinosa, E. Energetic, topological and electric field analyses of cation-cation nucleic acid interactions in Watson-Crick disposition. ChemPhysChem 2019, 20, 148–158.

(62) Frank-Kamenetskii, M. D.; Mirkin, S. M. Triplex DNA structures. Annu. Rev. Biochem. 1995, 64, 65–95.

(63) Jain, A.; Wang, G.; Vasquez, K. M. DNA triple helices: biological consequences and therapeutic potential. Biochimie 2008, 90, 1117–1130.

(64) Das, J.; Mukherjee, S.; Mitra, A.; Bhattacharyya, D. Non-Canonical base pairs and higher order structures in nucleic acids: Crystal structure database analysis. J. Biomol. Struct. Dyn. 2006, 24, 149–161.

(65) Lu, X.-J.; Olson, W. K. 3DNA: a versatile, integrated software system for the analysis, rebuilding and visualization of three-dimensional nucleic-acid structures. Nat. Protoc. 2008, 3, 1213.

(66) Bernier, C. R.; Petrov, A. S.; Waterbury, C. C.; Jett, J.; Li, F.; Freil, L. E.; Xiong, X.; Wang, L.; Migliozzi, B. L.; Hershkovits, E., et al. RiboVision suite for visualization and analysis of ribosomes. Farad. Discuss. 2014, 169, 195–207.

(67) Čech, P.; Svozil, D.; Hoksza, D. SETTER: web server for RNA structure comparison. Nucleic Acids Res. 2012, 40, W42–W48.

(68) Jmol: an open-source Java viewer for chemical structures in 3D. http://www.jmol.org/.

(69) Jones, R. A.; Steckelberg, A.-L.; Vicens, Q.; Szucs, M. J.; Akiyama, B. M.; Kieft, J. S. Different tertiary interactions create the same important 3-D features in a distinct flavivirus xrRNA. RNA 2021, 27, 54–65.

(70) Westhof, E.; Liang, S.; Tong, X.; Ding, X.; Zheng, L.; Dai, F. Unusual tertiary pairs in eukaryotic tRNA^*Ala*^. RNA 2020, 26, 1519–1529.

(71) Leontis, N. B.; Zirbel, C. L. Nonredundant 3D structure datasets for RNA knowledge extraction and benchmarking ; Springer, 2012; Vol. 27; pp 281–298.

(72) Ray, S. S.; Halder, S.; Kaypee, S.; Bhattacharyya, D. HD-RNAS: An automated hierarchical database of RNA structures. Front. Genet. 2012, 3, 1–10.

(73) Halder, A.; Roy, R.; Bhattacharyya, D.; Mitra, A. How does Mg^2+^ modulate the RNA folding mechanism: A case study of the G:C W:W Trans basepair. Biophys. J. 2017, 113, 277–289.

(74) Bhattacharya, S.; Jhunjhunwala, A.; Halder, A.; Bhattacharyya, D.; Mitra, A. Going beyond base-pairs: topology-based characterization of base-multiplets in RNA. RNA 2019, 25, 573–589.

(75) Gatti, C.; Macetti, G.; Boyd, R. J.; Matta, C. F. An electron density source-function study of DNA base pairs in their neutral and ionized ground states. J. Comp. Chem. 2018, 39, 1112–1128.

(76) Dennington, R.; Keith, T.; Millam, J. GaussView Version 5. Semichem Inc. Shawnee Mission KS 2009.

(77) Zhao, Y.; Schultz, N. E.; Truhlar, D. G. Design of density functionals by combining the method of constraint satisfaction with parametrization for thermochemistry, thermochemical kinetics, and noncovalent interactions. J. Chem. Theory Comput. 2006, 2, 364–382.

(78) Ditchfield, R.; Hehre, W. J.; Pople, J. A. Self-consistent molecular-orbital methods. IX. An extended gaussian-type basis for molecular-orbital studies of organic molecules. J. Chem. Phys. 1971, 54, 724–728.

(79) Sirianni, D. A.; Alenaizan, A.; Cheney, D. L.; Sherrill, C. D. Assessment of density functional methods for geometry optimization of bimolecular van der Waals complexes. J. Chem. Theo. Comput. 2018, 14, 3004–3013.

(80) Mardirossian, N.; Head-Gordon, M. How accurate are the Minnesota density functionals for noncovalent interactions, isomerization energies, thermochemistry, and barrier heights involving molecules composed of main-group elements? J. Chem. Theo. Comput. 2016, 12, 4303–4325.

(81) Halder, A.; Datta, A.; Bhattacharyya, D.; Mitra, A. Why does substitution of thymine by 6-ethynylpyridone increase the thermostability of DNA double helices? J. Phys. Chem. B 2014, 118, 6586–6596.

(82) Zhao, Y.; Truhlar, D. G. Density Functionals for Noncovalent Interaction Energies of Biological Importance. J. Chem. Theory Comput. 2007, 3, 289–300.

(83) Wang, J.; Gu, J.; Leszczynski, J. The electronic spectra and the H-bonding pattern of the sulfur and selenium substituted guanines. J. Comput. Chem. 2012, 33, 1587–1593.

(84) Banyasz, A.; Ketola, T.; Martínez-Fernández, L.; Improta, R.; Markovitsi, D. Adenine radicals generated in alternating AT duplexes by direct absorption of low-energy UV radiation. Faraday Discuss. 2018, 207, 181–197.

(85) Becke, A. D. Density-functional exchange-energy approximation with correct asymptotic behavior. Phys. Rev. A 1988, 38, 3098–3100.

(86) Lee, C.; Yang, W.; Parr, R. G. Development of the Colle-Salvetti correlation-energy formula into a functional of the electron density. Phys. Rev. B 1988, 37, 785–789.

(87) Miehlich, B.; Savin, A.; Stoll, H.; Preuss, H. Results obtained with the correlation energy density functionals of Becke and Lee, Yang and Parr. Chem. Phys. Lett. 1989, 157, 200–206.

(88) Yanai, T.; Tew, D. P.; Handy, N. C. A new hybrid exchange–correlation functional using the Coulomb-attenuating method (CAM-B3LYP). Chem. Phys. Lett. 2004, 393, 51–57.

(89) Grimme, S.; Antony, J.; Ehrlich, S.; Krieg, H. A consistent and accurate ab initio parametrization of density functional dispersion correction (DFT-D) for the 94 elements H-Pu. J. Chem. Phys. 2010, 132, 154104.

(90) Grimme, S.; Ehrlich, S.; Goerigk, L. Effect of the damping function in dispersion corrected density functional theory. J. Comput. Chem. 2011, 32, 1456–1465.

(91) Adamo, C.; Barone, V. Toward reliable density functional methods without adjustable parameters: The PBE0 model. J. Chem. Phys. 1999, 110, 6158–6170.

(92) Chai, J.-D.; Head-Gordon, M. Long-range corrected hybrid density functionals with damped atom–atom dispersion corrections. Phys. Chem. Chem. Phys. 2008, 10, 6615–6620.

(93) Humphrey, W.; Dalke, A.; Schulten, K. VMD – Visual molecular Dynamics. J. Mol. Graph. 1996, 14, 33–38.

(94) Frisch, M. J. et al. Gaussian 09 Revision C.01. Gaussian Inc. Wallingford CT 2009.

(95) Glendening, E. D.; Reed, A. E.; Carpenter, J. E.; Weinhold, F. NBO Version 3.1. 2004; Gaussian, Inc., Wallingford, CT, 2004.

(96) Bader, R. F. W., Ed. Atoms in molecules: A quantum theory, 1st ed.; Oxford Science: Oxford, 1990.

(97) Bader, R. F. W. A quantum theory of molecular structure and its applications. Chem. Rev. 1991, 91, 893–928.

(98) Todd A. Keith, T. G. AIMAll (Version 11.12.19). 2012; Overland Park KS, USA.

(99) Møller, C.; Plesset, M. S. Note on an Approximation Treatment for Many-Electron Systems. Phys. Rev. 1934, 46, 618–622.

(100) Becke, A. D. Density-functional thermochemistry. V. Systematic optimization of exchange-correlation functionals. J. Chem. Phys. 1997, 107, 8554–8560.

(101) Grimme, S. Semiempirical GGA-type density functional constructed with a long-range dispersion correction. J. Comput. Chem. 2006, 27, 1787–1799.

(102) Grimme, S. Semiempirical hybrid density functional with perturbative second-order correlation. J. Chem. Phys. 2006, 124, 034108.

(103) Antony, J.; Brüske, B.; Grimme, S. Cooperativity in noncovalent interactions of biologically relevant molecules. Phys. Chem. Chem. Phys. 2009, 11, 8440–8447.

(104) Boys, S. F.; Bernardi, F. The calculation of small molecular interactions by the differences of separate total energies. Some procedures with reduced errors. Mol. Phys. 1970, 19, 553–566.

(105) Cossi, M.; Rega, N.; Scalmani, G.; Barone, V. Energies, structures, and electronic properties of molecules in solution with the C-PCM solvation model. J. Comput. Chem. 2003, 24, 669–681.

(106) Saha, S.; Sastry, G. N. Quantifying cooperativity in water clusters: An attempt towards obtaining a generalised equation. Mol. Phys. 2015, 113, 3031–3041.

(107) Oliva, R.; Cavallo, L.; Tramontano, A. Accurate energies of hydrogen bonded nucleic acid base pairs and triplets in tRNA tertiary interactions. Nucleic Acids Res. 2006, 34, 865–879.

(108) Preethi, S.; Sharma, P.; Mitra, A. Higher order structures involving post transcriptionally modified nucleobases in RNA. RSC Adv. 2017, 7, 35694–35703.

(109) Oliva, R.; Tramontano, A.; Cavallo, L. Mg^2+^ binding and archaeosine modification stabilize the G15–C48 Levitt base pair in tRNAs. RNA 2007, 13, 1427–1436.

(110) Levitt, M. Detailed molecular model for transfer ribonucleic acid. Nature 1969, 224, 759–763.

(111) Halder, A.; Data, D.; Seelam, P. P.; Bhattacharyya, D.; Mitra, A. Estimating strengths of individual hydrogen bonds in RNA base pairs: Toward a consensus between different computational approaches. ACS Omega 2019, 4, 7354–7368.

(112) Miao, Z. et al. RNA-Puzzles Round II: assessment of RNA structure prediction programs applied to three large RNA structures. RNA 2015, 21, 1066–1084.

(113) Garg, A.; Heinemann, U. A novel form of RNA double helix based on G·U and C·A+ wobble base pairing. RNA 2018, 24, 209–218.

(114) Frye, M.; Harada, B. T.; Behm, M.; He, C. RNA modifications modulate gene expression during development. Science 2018, 361, 1346–1349.

(115) Song, J.; Yi, C. Chemical modifications to RNA: A new layer of gene expression regulation. ACS Chem. Biol. 2017, 12, 316–325.

(116) Barbieri, I.; Kouzarides, T. Role of RNA modifications in cancer. Nat. Rev. Cancer 2020, 20, 1–20.

