## Supporting Information for "Occurrences of protonated base triples in RNA are determined by their cooperative binding energies and specific functional requirements"

**Total number of possible base pairing interactions between two RNA nucleobases:**

- All unique base-base combinations possible in an RNA base pair.
  - 4 homo dimers
  - 6 hetero dimers

| Base 1 \ Base 2 | A | U | G | C |
|---|---|---|---|---|
| **A** | AA | AU | AG | AC |
| **U** | | UU | UG | UC |
| **G** | | | GG | GC |
| **C** | | | | CC |

- All unique interacting edge combinations possible for homo dimers = 6
- Each combination could be either *cis* or *trans*
- Hence, total possibilities = 6X2 = 12

| Edge 1 \ Edge 2 | W | H | S |
|---|---|---|---|
| W | WW | WH | WS |
| H | | HH | HS |
| S | | | SS |

- Total number of base pairing geometries possible for homo dimers = 4X12 = 48

- All unique interacting edge combinations possible for hetero dimers = 9
- Each combination could be either *cis* or *trans*
- Hence, total possibilities = 9X2 = 18

| Edge 1 \ Edge 2 | W | H | S |
|---|---|---|---|
| **W** | WW | WH | WS |
| **H** | HW | HH | HS |
| **S** | SW | SH | SS |

- Total number of base pairing geometries possible for hetero dimers = 6X18 = 108

- **Hence total number of possible base pairing interactions = 48 + 108 = 156**

Out of these 156 possible base pairing geometries, 118 distinct base-pair varieties (in terms of both base combination and geometry), observed in available RNA structures, are reported in the NDB RNA Base pair catalog: `http://ndbserver.rutgers.edu/ndbmodule/services/BPCatalog/bpCatalog.html`

**Annotation of a base pairing interaction:** As per Leontis and Westhof (LW) nomenclature scheme, base pairing interaction between edge X of base M and edge Y of base N in cis orientation is represented as MN cXY. However, the nomenclature used in this thesis is slightly different. The same base pair is represented as M:N X:Y Cis. This sort of representation is also intuitive and at the same time, due to the presence of the text delimiter (:), is more convenient for parsing of metadata files, especially when modified nucleobases (represented by more than one character, e.g. 7MG, 2MA, etc) are involved. If the inter-base hydrogen bonds in a particular base pair involve any of the sugar atoms (O2′, O3′, etc) I add a prefix 'r' with the base, e.g., G:rC W:S Cis.

Table S1: Occurrence frequencies of protonated base triples in the 'larger dataset'

(i) Part 1

| Base Triple | Category | Count |
|---|---|---|
| A(CC) +:WC/H:WT | Type I | 318 |
| C(AA) W:+C/S:SC | Type II | 174 |
| G(CC) H:+T/W:WC | Type II | 155 |
| G(AC) H:zC/W:WC | Type II | 106 |
| A(GA) +:HC/S:SC | Type I | 28 |
| G(AU) H:+T/W:WC | Type II | 19 |
| G(CA) W:+C/S:WT | Type II | 19 |
| A(CA) +:WC/S:SC | Type I | 16 |
| G(CA) W:+C/S:ST | Type II | 14 |
| C(GG) +:WC/W:WC | Type I | 9 |
| C(CG) W:+C/W:WC | Type II | 7 |
| G(AC) H:+T/W:WC | Type II | 6 |
| G(GU) z:HT/W:WC | Type I | 6 |
| G(CC) H:+C/W:WC | Type II | 5 |
| G(CU) H:+T/W:WC | Type II | 5 |
| G(CC) S:+T/W:WC | Type II | 5 |
| G(CC) z:WC/W:WC | Type I | 5 |
| C(GU) +:WC/H:SC | Type I | 4 |
| C(GA) +:WC/S:WC | Type I | 4 |
| G(CC) W:+C/W:WC | Type II | 3 |
| U(CA) W:+C/W:WC | Type II | 3 |
| A(GU) +:HC/H:WT | Type I | 2 |
| C(GU) +:HC/W:WC | Type I | 2 |
| C(CG) +:WC/W:WC | Type I | 2 |
| C(UG) +:WC/W:WC | Type I | 2 |
| G(AG) H:+C/S:ST | Type II | 2 |
| G(GG) H:+T/S:ST | Type II | 2 |
| C(CG) S:+T/W:WC | Type II | 2 |

(i) Part 2

| Base Triple | Category | Count |
|---|---|---|
| G(CC) S:+T/W:WT | Type II | 2 |
| G(CG) W:+C/H:WC | Type II | 2 |
| G(CG) W:+C/H:WT | Type II | 2 |
| C(AC) W:+C/S:HC | Type II | 2 |
| G(CG) W:+C/S:SC | Type II | 2 |
| A(GU) +:HC/S:SC | Type I | 1 |
| C(GA) +:HT/S:HC | Type I | 1 |
| C(GC) +:WC/W:WC | Type I | 1 |
| A(CA) +:WC/H:SC | Type I | 1 |
| C(GA) +:WC/S:SC | Type I | 1 |
| C(GG) +:WC/S:SC | Type I | 1 |
| C(GC) +:WC/S:WC | Type I | 1 |
| C(GG) +:WC/S:WC | Type I | 1 |
| C(GA) +:WC/S:WT | Type I | 1 |
| C(GG) +:WC/S:WT | Type I | 1 |
| C(CA) +:WC/W:HT | Type I | 1 |
| C(GA) +:WC/W:WC | Type I | 1 |
| C(CG) +:WT/H:HT | Type I | 1 |
| C(UA) +:WT/W:HT | Type I | 1 |
| G(AC) H:+C/W:SC | Type II | 1 |
| G(CC) H:+C/W:SC | Type II | 1 |
| G(GC) H:+T/W:WC | Type II | 1 |
| U(CA) W:+C/S:SC | Type II | 1 |
| C(CG) W:+C/S:WT | Type II | 1 |
| C(AA) W:+C/W:WC | Type II | 1 |
| G(CA) W:+C/W:WC | Type II | 1 |
| G(CU) W:+C/W:WC | Type II | 1 |
| U(CA) W:+T/S:WC | Type II | 1 |

**Basic statistics:**

Total types : 56; Total instances : 958

Type I : 26 types (412 instances), Type II : 30 types (546 instances)

Table S2: Occurrence frequencies of protonated base triples in the NDB dataset.

| Base Triple | Category | Count |
|---|---|---|
| A(CC) +:WC/H:WT | Type I | 9 |
| G(CC) H:+T/W:WC | Type II | 9 |
| G(AU) H:+T/W:WC | Type II | 7 |
| C(AA) W:+C/S:SC | Type II | 6 |
| G(CC) H:+C/W:WC | Type II | 5 |
| G(CC) +:WC/W:WC | Type I | 4 |
| A(CA) +:WC/S:SC | Type I | 3 |
| A(GA) +:HC/S:SC | Type I | 3 |
| C(CG) W:+C/W:WC | Type II | 3 |
| G(AC) H:+T/W:WC | Type II | 3 |
| A(CU) +:WC/H:WC | Type I | 1 |
| A(GU) +:HC/S:SC | Type I | 1 |
| C(CG) +:WT/H:HT | Type I | 1 |
| C(GG) +:HC/W:ST | Type I | 1 |
| C(GG) +:WC/W:WC | Type I | 1 |
| G(AC) H:+C/W:WC | Type II | 1 |
| G(CA) W:+C/S:WT | Type II | 1 |

**Basic statistics:**
Total types : 17; Total instances : 59
Type I : 9 types (24 instances), Type II : 8 types (35 instances)

Table S3: Occurrence frequencies of protonated base triples in the HDRNAS dataset.

| Base Triple | Category | Count |
|---|---|---|
| A(CC) +:WC/H:WT | Type I | 8 |
| C(AA) W:+C/S:SC | Type II | 6 |
| G(AU) H:+T/W:WC | Type II | 5 |
| G(CC) H:+T/W:WC | Type II | 5 |
| G(CC) H:+C/W:WC | Type II | 4 |
| G(AC) H:+C/W:WC | Type II | 3 |
| A(GA) +:HC/S:SC | Type I | 1 |
| A(GU) +:HC/S:SC | Type I | 1 |
| C(AC) W:+C/S:HC | Type II | 1 |
| G(AC) H:+T/W:WC | Type II | 1 |
| G(CA) W:+C/S:ST | Type II | 1 |

**Basic statistics:**
Total types : 11; Total instances : 36
Type I : 3 types (10 instances), Type II : 8 types (26 instances)

Table S4: Instances of protonated base triples that are selected for QM calculations. Each triple is composed of two base pairs (one protonated and one neutral). E-values corresponding to the constituent base pairs are reported.

| System | Protonated Base Triple | PDB | Chain | Base Pair 1 | E-value | Base Pair 2 | E-value |
|---|---|---|---|---|---|---|---|
| 1 | A(CC) +:WC/H:WT | 2ZJQ | X | 2429A:2046C +:W Cis | 0.85 | 2429A:2480C H:W Trans | 1.7 |
| 2 | G(CC) H:+T/W:WC | 4W2G | B (bundle 4) | 67G:17C W:W Cis | 0.51 | 67G:109C H:+ Trans | 0.91 |
| 3 | C(AA) W:+C/S:SC | 4WZO | A (bundle 2) | 964C:953A W:+ Cis | 0.8 | 964C:2273A S:S Cis | 1.67 |
| 4 | G(AU) H:+T/W:WC | 1IL2 | D | 1922G:1913PSU W:W Cis | 0.18 | 1922G:1946A H:+ Trans | 0.78 |
| 5 | G(CC) H:+C/W:WC | 4PLX | C | 71G:12C H:+ Cis | 0.38 | 71G:42C W:W Cis | 0.31 |
| 6 | G(AC) H:zC/W:WC | 2O44 | A | 2035G:2596C W:W Cis | 0.96 | 2035G:2034A H:z Cis | 0.52 |
| 7 | A(GA) +:HC/S:SC | 4U56 | D (bundle 3) | 407A:390G +:H Cis | 0.67 | 407A:1671A S:S Cis | 1.07 |
| 8 | G(AC) H:+T/W:WC | 4U56 | D (bundle 2) | 2396G:2984C W:W Cis | 0.39 | 2396G:2946A H:+ Trans | 1.45 |
| 9 | G(CC) z:WC/W:WC | 1ET4 | A | 226G:219C W:W Cis | 0.22 | 226G:324C z:W Cis | 0.4 |
| 10 | A(CA) +:WC/S:SC | 4U52 | I (bundle 4) | 2107A:1941C +:W Cis | 0.25 | 2107A:3344A S:S Cis | 1.5 |
| 11[a] | C(CG) W:+C/W:WC | 4V5A | A (bundle 2) | 2803C:2793G W:W Cis | 1.66 | 2803C:2794C W:+ Cis | 1.29 |
| 12 | A(GU) +:HC/S:SC | 2A64 | A | 385A:47G +:H Cis | 0.69 | 385A:400U S:S Cis | 0.88 |
| 13 | G(CA) W:+C/S:WT | 2OTJ | 0 | 940G:1032A S:W Trans | 1.53 | 940G:1026C W:+ Cis | 0.52 |
| 14 | G(CA) W:+C/S:ST | 4U25 | A (bundle 4) | 333G:318C W:+ Cis | 0.91 | 333G:300A S:S Trans | 0.82 |
| 15[a] | C(GG) +:WC/W:WC | 4U52 | A (bundle 1) | 276C:281G W:W Cis | 1.48 | 276C:279G +:W Cis | 0.83 |
| 16 | C(AC) W:+C/S:HC | 2ZJR | X | 937C:865A W:+ Cis | 0.36 | 937C:939C S:H Cis | 0.84 |
| 17 | C(CG) +:WT/H:HT | 3EGZ | B | 57C:20C +:W Trans | 0.44 | 57C:21G H:H Trans | 1.49 |
| 18 | A(CU) +:WC/H:WC | 4W23 | 2 | 650A:649U H:W Cis | 1.42 | 650A:24C +:W Cis | 0.13 |
| 19[a] | C(GG) +:HC/W:ST | 4W21 | 5 | 4337 C:4373G +:H Cis | 0.95 | 4337C:4228G W:S Trans | 1.35 |
| 20 | G(GU) z:HT/W:WC | 4WQ1 | A (bundle 1) | 1058G:1202G z:H Trans | 1.11 | 1058G:1199U A W:W Cis | 0.79 |

[a] These systems are bifurcated triples, i.e., one single edge of the central base participates in base pairing interactions with two terminal bases.

Table S5: Comparison of the optimized geometries of two protonated base triples, G(CC) z:WC/W:WC (Type I) and G(CC) H:+T/W:WC (Type II) obtained using six different DFT functionals. These functionals include (a) the hybrid meta-GGA functional M05-2X that is parameterized for middle range dispersion correstions, (b) B3LYP[1–3] which is arguably the most widely used fucntional for nucleobases and related systems, (c) its long range corrected version CAM-B3LYP,[4] (d) its dispersion corrected version B3LYP-D3(BJ), where dispersion is added explicitly using Grimme's method ($3^{rd}$ order) with Becke-Johnson damping,[5,6] (e) PBE0 where all the parameters (except those in the underlying local spin-density approximation) are physical constants[7] and (f) $\omega$B97-XD[8] which is recommended for DNA triples by Grimme and co-workers.[9] The basis set used is 6-31++G(2d,2p). Another calculation is done with M05-2X functional and a larger basis set 6-311++G(2df,2pd). RMSD(1) and RMSD(2) values are calculated with respect to the optimized geometry obtained at M05-2X/6-31++G(2d,2p) level and reported in Å.

| Level of theory | G(CC) z:WC/W:WC | | G(CC) H:+T/W:WC | |
|---|---|---|---|---|
| | RMSD(1) | RMSD(2) | RMSD(1) | RMSD(2) |
| (a) M05-2X/6-31++G(2d,2p) | 0.00 | 0.00 | 0.00 | 0.00 |
| (b) B3LYP/6-31++G(2d,2p) | 0.03 | 0.05 | 0.10 | 0.13 |
| (c) CAM-B3LYP/6-31++G(2d,2p) | 0.03 | 0.04 | 0.10 | 0.12 |
| (d) B3LYP-D3(BJ)/6-31++G(2d,2p) | 0.10 | 0.11 | 0.10 | 0.13 |
| (e) PBE0/6-31++G(2d,2p) | 0.06 | 0.09 | 0.10 | 0.13 |
| (f) $\omega$B97-XD/6-31++G(2d,2p) | 0.12 | 0.26 | 0.20 | 0.26 |
| (g) M05-2X/6-311++G(2df,2pd) | 0.01 | 0.01 | 0.07 | 0.09 |

Table S6: Protonated triples that occur recurrently in different ribosomal RNAs and ribozymes.

| System | Geometry | Type of RNA | Species | Constituent BPs | Structural context of occurrence |
|---|---|---|---|---|---|
| 1 | A(CC) +:WC/H:WT | 23S rRNA | *T. thermophilus* | 2450A:2501C H:W Trans; 2450A:2063C +:WC Trans | Multi junction loop in Domain V (connects 5 helices) |
| | | 23S rRNA | *D. radiodurans* | 2429A:2046C +:W Cis; 2429A:2480C H:W Trans | |
| | | 23S rRNA | *E. coli* | 2450A:2501C H:W Trans; 2450A2063C +:W Cis | |
| | | 23S rRNA | *H. marismortui* | 2485A:2536C H:W Trans; 2485A:2104C +:W Cis | |
| | | 25S rRNA | *S. cerevisiae* | 2819A:2870C H:W Trans; 2819A:2405C +:W Cis | |
| | | 28S rRNA | *P. falciparum* | 3178A:2698C +:W Cis; 3178A:3229C H:W Trans | |
| | | Mitochondrial 16S rRNA | *H. sapiens* | 2937A:2726C +:W Cis; 2937A:2988C H:W Trans | Multi junction loop (connects 3 helices) |
| | | Mitochondrial 16S rRNA | *S. scrofa* | 1271A:1056C +:W Cis; 1271A:1322C H:W Trans | |
| | | 23S rRNA | *H. marismortui* | 1005A:963C +:W Cis; 1005A:959C H:W Trans | Internal loop in Domain II |
| 2 | G(CC) H:+T/W:WC | 5S rRNA | *T. thermophilus* | 67G:17C W:W Cis; 67G:109C H:+ Trans | Multi junction loop (connects 3 helices) |
| | | 5S rRNA | *H. marismortui* | 66G:15C W:W Cis; 66G:113C H:+ Trans | |
| | | 23S rRNA | *S. cerevisiae* | 1513G:1505C W:W Cis; 1513G:1502C H:+ Trans | Internal loop in Domain III |
| | | 28S rRNA | *P. falciparum* | 1662G:1654C W:W Cis; 1662G:1651C H:+ Trans | |
| | | HDV ribozyme | Hepatitis delta virus | 161G:144C W:W Cis; 161G:141C H:+ Trans | Junction of P4 and P1.1 |
| 3 | C(AA) W:+C/S:SC | 23S rRNA | *T. thermophilus* | 964C:953A W:+ Cis; 964C:2273A S:S Cis | Loop-loop interaction between one internal loop (of Domain II) and one hairpin loop (of Domain V) |
| | | 23S rRNA | *D. radiodurans* | 975C:964A W:+ Cis; 975C:2252A S:S Cis | |
| | | 18S rRNA | *S. cerevisiae* | 625C:974A W:+ Cis; 625C:939A S:S Cis | Loop-loop interaction between one bulge loop and a GAAA tetra loop of Domain C |
| | | 18S rRNA | *P. falciparum* | 632C:1043A W:+ Cis; 632C:1008A S:S Cis | |
| | | 18S rRNA | *S. scrofa* | 674C:1031A W:+ Cis; 674C:996A S:S Cis | |
| | | mitochondrial 16S rRNA | *H. sapiens* | 2538C:2512A W:+ Cis; 2538C:2633A S:S Cis | Similar loop-loop interaction between one bulge loop and a GAAA tetra loop |

**Table S6 – continued from previous page**

| System | Geometry | Type of RNA | Species | Constituent BPs | Structural context of occurrence |
|---|---|---|---|---|---|
| | | mitochondrial 16S rRNA | *S. scrofa* | 868C:842A W:+ Cis; 868C963A S:S Cis | |
| 7 | A(GA) +:HC/S:SC | 18S rRNA | *S. cerevisiae* | 407A:390G +:H Cis; 407A:1671A S:S Cis | Loop-loop interaction between one internal loop in Domain 5′ with another internal loop in Domain 3′m |
| | | | *P. falciparum* | 413A:396G +:H Cis; 413A:1968A S:S Cis | |
| | | | *S. scrofa* | 455A:438G +:H Cis; 455A:1735A S:S Cis | |

Table S7: Protonated base triples found in different ribosomal RNAs. These occurrence instances are not recurrent.

| System | Geometry | Type of RNA | Species | Structural context of occurrence |
|---|---|---|---|---|
| 6 | G(AC) H:zC/W:WC | 23S rRNA | *T. thermophilus* | Part of a bulge loop in Domain 0 |
| 10 | A(CA) +:WC/S:SC | Mitochondrial 16S rRNA | *H. sapiens* | Loop-loop interaction between a junction loop to a hairpin loop |
| | | 18S rRNA | *S. scrofa* | Loop-loop interaction between a junction loop to a hairpin loop |
| | | 25S rRNA | *S. cerevisiae* | Loop-loop interaction between one bulge loop of Domain VI and another bulge loop of Domain IV |
| 11 | C(CG) W:+C/W:WC | 28S rRNA | *S. scrofa* | Within double helical regions |
| | | SRP RNA | *P. horikoshii* | Within double helical regions |
| 13 | G(CA) W:+C/S:WT | 23S rRNA | *H. marismortui* | Part of an internal loop in Domain II |
| 14 | G(CA) W:+C/S:ST | 23S rRNA | *H. marismortui* | Part of an internal loop in Domain II |
| 15 | C(GG) +:WC/W:WC | 28S rRNA | *P. falciparum* | Part of a multi junction loop (connects 3 helices) |
| 16 | C(AC) W:+C/S:HC | 23S rRNA | *D. radiodurans* | Part of an internal loop |
| 18 | A(CU) +:WC/H:WC | 18S rRNA | *S. scrofa* | Within double helical regions |
| 19 | C(GG) +:HC/W:ST | 28S rRNA | *S. scrofa* | Part of a multi junction loop (connects 3 helices) |
| 20 | G(GU) z:HT/W:WC | 16S rRNA | *T. thermophilus* | Multi Junction loop |

Table S8: Protonated base triples that are found in synthetic RNA constructs.

| System | Gemetry | Type of RNA | Structural context of occurrence |
|---|---|---|---|
| 2 | G(CC) H:+T/W:WC | Viral RNA Pseudoknot | Part of a pseudoknot |
| 5 | G(CC) H:+C/W:WC | tRNA CCA-acceptor | Tertiary contact, connecting two different chains |
| 9 | G(CC) z:WC/W:WC | Vitamin B12 binding RNA aptamer | Tertiary contact, connecting two different chains |
| 11 | C(CG) W:+C/W:WC | ds RNA | Within double helical stretches |
| 12 | A(GU) +:HC/S:SC | ribonuclease P RNA | Part of a 4 way junction loop |
| 17 | C(CG) +:WT/H:HT | Tetracycline aptamer and artificial riboswitch | Mediates helix-helix interaction |

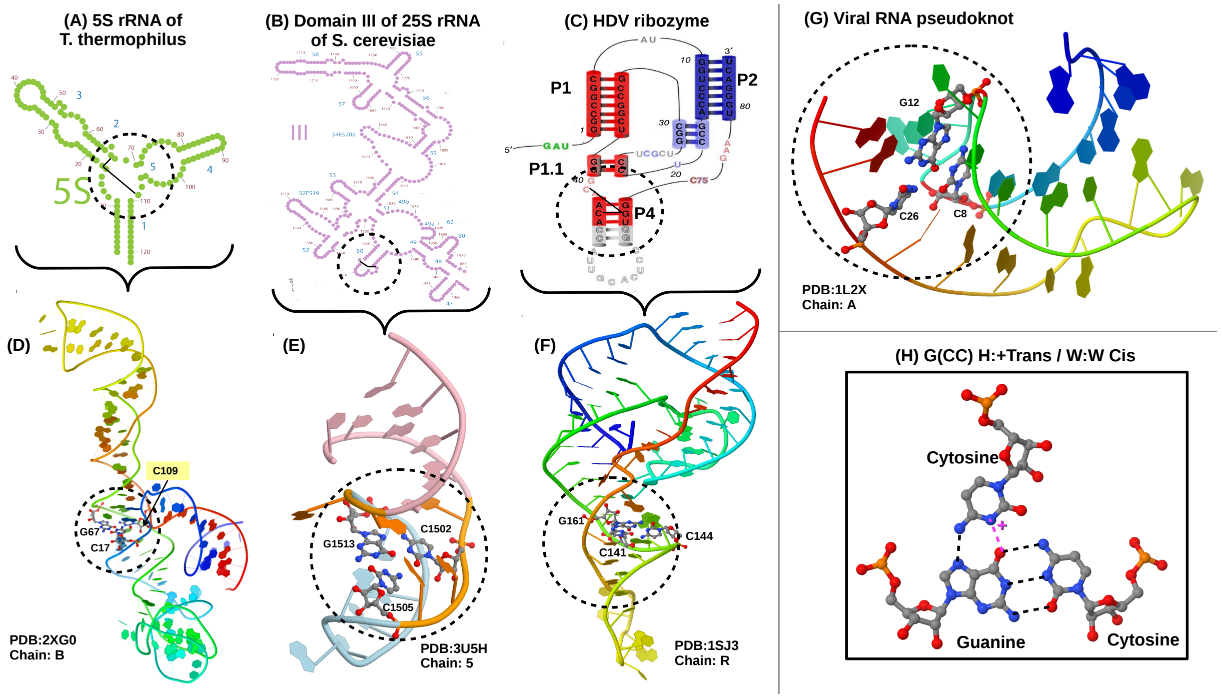


Figure S1: **Context of occurrences of G(CC) H:+ Trans/W:W Cis triple (*System 2*).** 2D representation of (A) the 5S rRNA, (B) Domain III of 25S rRNA of *S. cerevisiae* and (C) HDV ribozyme are shown and the structural motifs that contain the base triple (*System 2*) are encircled. 3D structures corresponding to these motifs shown in (B), (C) and (D) are illustrated in (D), (E) and (F), respectively. (G) 3D structure of the viral RNA pseudoknot containing *System 02*. (H) Geometry of the base triple. Inter-base hydrogen bonds are shown in broken lines (magenta color represents the proton mediated hydrogen bond).

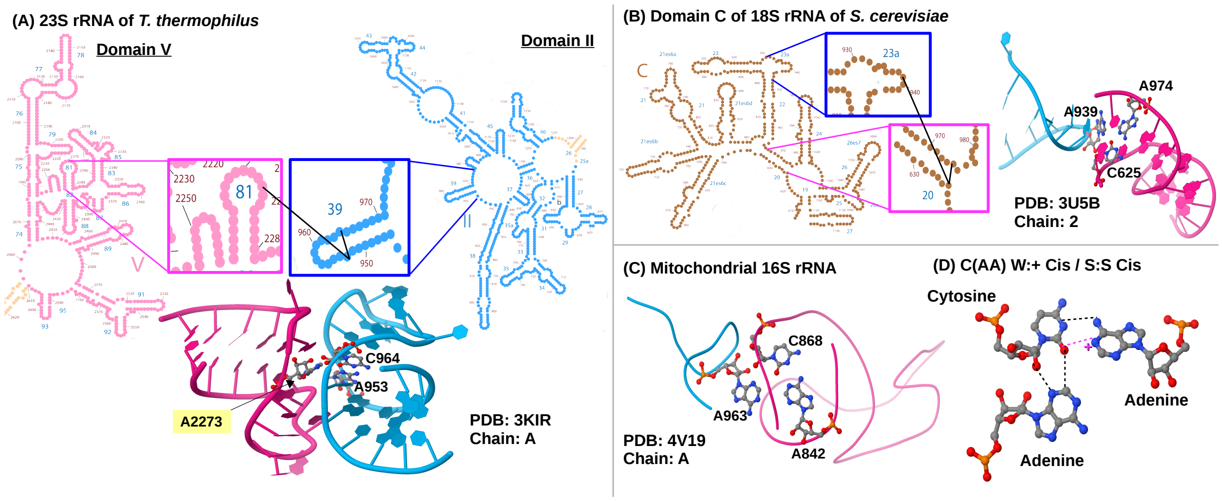


Figure S2: **Context of occurrences of C(AA) W:+ Cis/S:S Cis triple (*System 3*).** (A) 2D and 3D illustrations of a loop loop interaction (mediated by *System 3*) between a hairpin loop of Domain V (pink color) and an internal loop in Domain II (blue color) of 23S rRNA of *T. thermophilus*. The black lines represent the base-base contacts in *System 3*. (B) 2D and 3D illustrations of a loop loop interaction (mediated by *System 3*) between a bulge loop and a GNRA tetra loop (GAAA in this case) in Domain C of 28S rRNA of *S. cerevisiae*. (C) Similar loop-loop interaction mediated by System 3 (as illustrated in (B)) is also observed in mitochondrial 16S rRNA of mammals. (D) Geometry of the base triple. Inter-base hydrogen bonds are shown in broken lines (magenta color represents the proton mediated hydrogen bond).

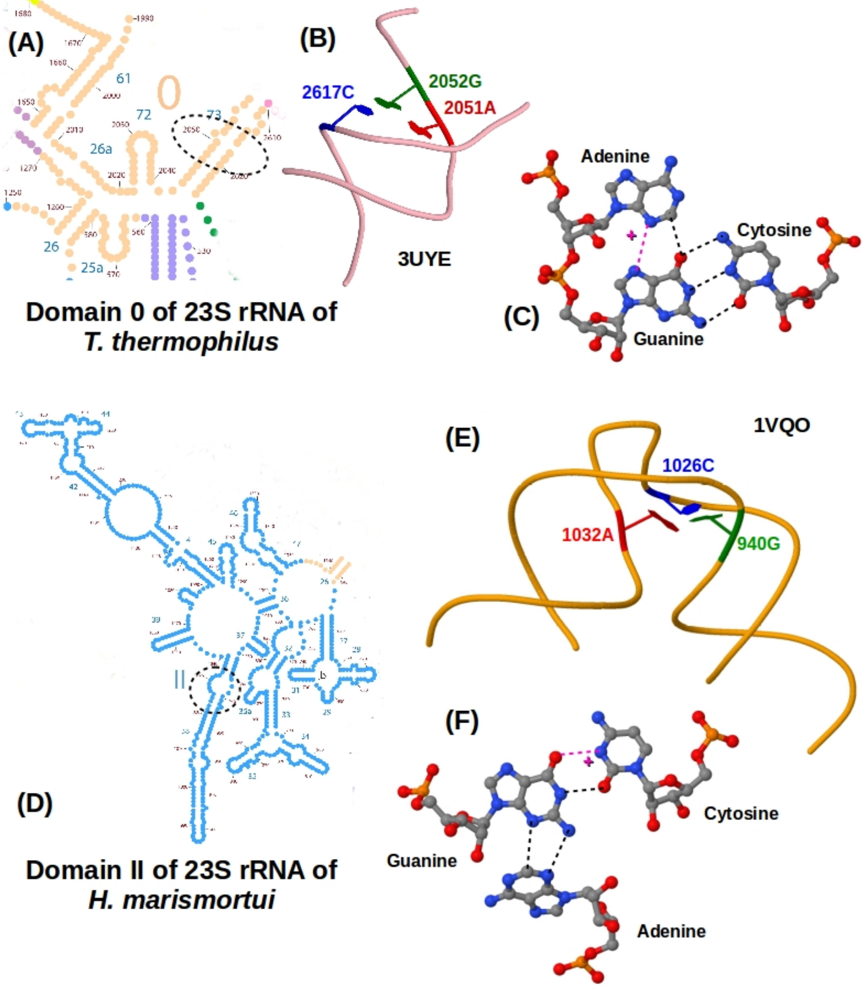


Figure S3: Context of occurrences of the important protonated triples that are detected in different rRNAs but the occurrences are not recurrent. **System 6 (G(AC) H:ZC/W:WC):** (A) 2D representation of Domain 0 of 23S rRNA of *T. thermophilus*. The bulge loop containing System 6 has been encircled. (B) 3D structure of the bulge loop. (C) Geometry of the base triple. Inter-base hydrogen bonds are shown in broken lines (magenta color represents the proton mediated hydrogen bond). **System 14 (G(CA) W:+C/S:ST):**(D) 2D representation of the Domain II of 23S rRNA of *H. marismortui*. The asymmetric internal loop containing System 14 has been encircled. (E) 3D structure of the internal loop. (F) Geometry of the base triple. Inter-base hydrogen bonds are shown in broken lines (magenta color represents the proton mediated hydrogen bond).

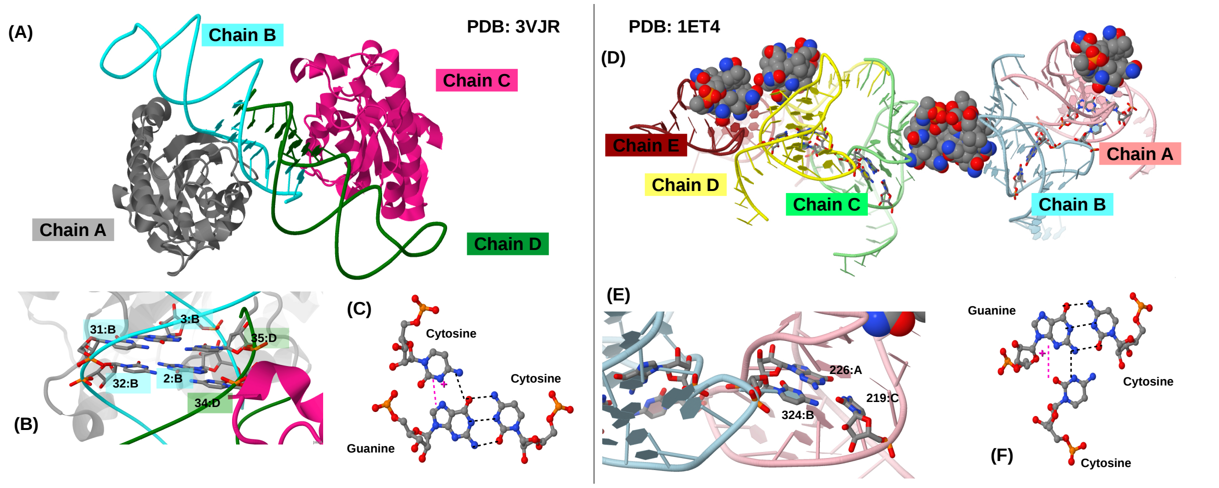


Figure S4: Context of occurrences of the important protonated triples found in synthetic RNA constructs. **System 5 (G(CC) H:+C/W:WC) :** (A) 3D structure of tRNA CCA-acceptor. System 5 connects chain B and chain D. (B) An enlarged view of the region containing two consecutively stacked G(CC) H:+C/W:WC triple. (C) Geometry of the base triples. Inter-base hydrogen bonds are shown in broken lines (magenta color represents the proton mediated hydrogen bond). **System 9 (G(CC) z:WC/W:WC):** (D) 3D structure of the Vitamin B12 binding RNA aptamer. The Vitamin B12 molecules are shown in spacefill model. Four different occurrences of System 9 are shown in stick model with CPK color scheme. (E) An enlarged view of the region containing System 9 between chain A and chain B. (F) Geometry of the base triple. Inter-base hydrogen bonds are shown in broken lines (magenta color represents the proton mediated hydrogen bond).

Table S9: Interaction energy ($\triangle$E) of the protonated triples calculated using M05-2X and $\omega$B97XD functionals with 6-311++G(2df,2pd) basis set. $\triangle$E values are further corrected for deformation energy ($E^{def}$) and BSSE ($E^{BSSE}$) to obtain the final interaction energy $E^{int}$ ($= \triangle E + E^{def} + E^{BSSE}$). CE represents the co-operative energy corresponding to the base triples. All the energy values are reported in kcal/mol.

| Sys. | Nomenclature of the | Category | M05-2X | | | | | $\omega$B97XD | | | | |
|---|---|---|---|---|---|---|---|---|---|---|---|---|
| No. | base triple | | $\triangle$E | $E^{def}$ | $E^{BSSE}$ | $E^{int}$ | CE | $\triangle$E | $E^{def}$ | $E^{BSSE}$ | $E^{int}$ | CE |
| 1 | A(CC) +:WC/H:WT | Type I | -76.0 | 6.6 | 1.3 | -68.1 | 0.9 | -79.1 | 6.7 | 1.3 | -71.1 | 0.9 |
| 2 | G(CC) H:+T/W:WC | Type II | -81.4 | 8.5 | 1.5 | -71.4 | -6.1 | -85.4 | 8.4 | 1.5 | -75.5 | -6.1 |
| 3 | C(AA) W:+C/S:SC | Type II | -65.7 | 11.5 | 2.0 | -52.2 | 3.5 | -71.1 | 10.4 | 1.9 | -58.8 | 3.5 |
| 4 | G(AU) H:+T/W:WC | Type II | -69.0 | 5.0 | 1.2 | -62.8 | 0.9 | -72.3 | 5.0 | 1.3 | -66.0 | 0.9 |
| 5 | G(CC) H:+C/W:WC | Type II | -82.2 | 10.0 | 1.4 | -70.8 | -3.6 | -86.2 | 9.9 | 1.4 | -74.9 | -3.6 |
| 6 | G(AC) H:zC/W:WC | Type II | -71.1 | 7.1 | 1.3 | -62.7 | -3.7 | -75.2 | 7.1 | 1.3 | -66.9 | -3.7 |
| 7 | A(GA) +:HC/S:SC | Type I | -63.8 | 7.4 | 1.8 | -54.6 | 0.6 | -69.1 | 6.8 | 1.8 | -60.6 | 0.6 |
| 8 | G(AC) H:+T/W:WC | Type II | -77.7 | 7.4 | 1.3 | -69.0 | -3.1 | -81.9 | 7.4 | 1.3 | -73.1 | -3.2 |
| 9 | G(CC) z:WC/W:WC | Type I | -89.9 | 6.7 | 1.4 | -81.8 | 5.4 | -94.0 | 6.8 | 1.4 | -85.8 | 5.4 |
| 10 | A(CA) +:WC/S:SC | Type I | -67.5 | 8.3 | 2.4 | -56.8 | 1.1 | -73.2 | 7.4 | 2.3 | -63.5 | 1.2 |
| 11 | C(CG) W:+C/W:WC | Type II | -67.8 | 5.8 | 2.7 | -59.3 | 3.8 | -72.9 | 5.7 | 2.5 | -64.7 | 3.5 |
| 12 | A(GU) +:HC/S:SC | Type I | -60.1 | 3.8 | 2.0 | -54.2 | 0.1 | -65.3 | 2.9 | 2.0 | -60.4 | 0.1 |
| 13 | G(CA) W:+C/S:WT | Type II | -70.0 | 16.7 | 1.7 | -51.6 | -1.9 | -72.7 | 16.6 | 1.6 | -54.5 | -1.9 |
| 14 | G(CA) W:+C/S:ST | Type II | -62.6 | 16.1 | 1.1 | -45.3 | -2.2 | -63.9 | 16.2 | 1.1 | -46.7 | -2.2 |
| 18 | A(CU) +:WC/H:WC | Type I | -65.1 | 4.4 | 1.2 | -59.4 | 1.2 | -68.7 | 4.5 | 1.3 | -62.9 | 1.2 |
| 20 | G(GU) z:HT/W:WC | Type I | -69.6 | 5.3 | 1.2 | -63.1 | 3.2 | -72.6 | 5.5 | 1.3 | -65.9 | 3.0 |

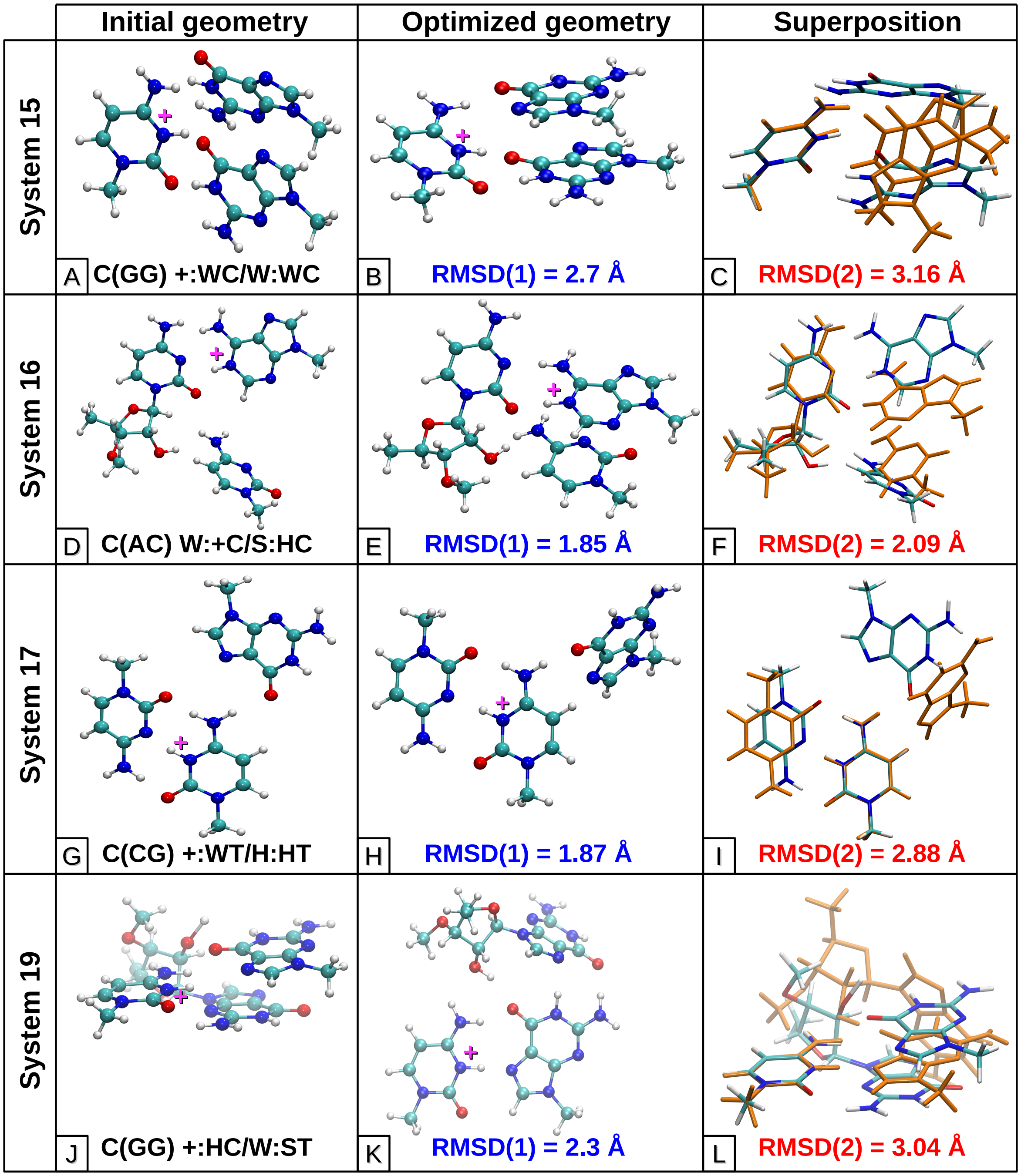


Figure S5: Initial (crystal geometry) and final optimized geometries (at M05-2X/6-31++G(2d,2p) level) of system 15, 16, 17 and 19. Final optimized geometries of these triples are significantly deviated from their respective crystal geometries, as characterized by RMSD(1) and RMSD(2) values. Initial and final geometries are superposed to each other with respect to the central base. The initial geometry is represented in orange color and the final geometry is represented in CPK color scheme. In general geometry optimization of all the systems were carried out in their respective internal coordinate spaces. However for these systems we carried out geometry optimization in Cartesian coordinate space as well, to be sure about the deviation from their respective native geometries.

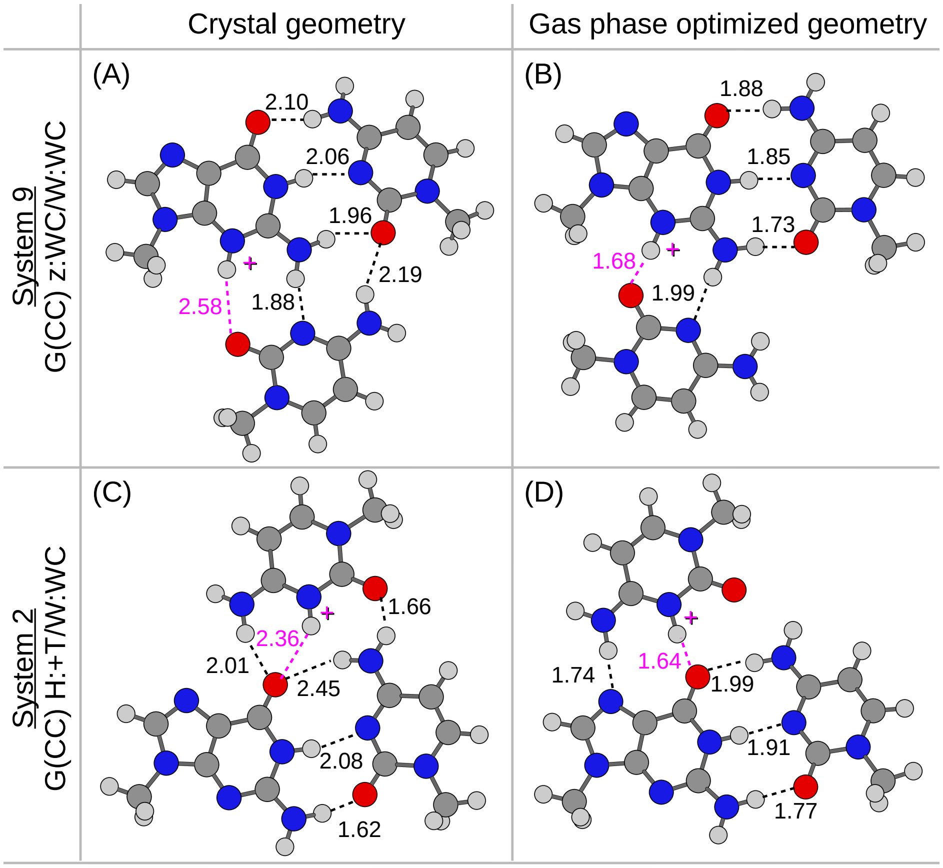


Figure S6: In some base triples, even though the overall geometries remain unchanged, inter-base hydrogen bonding pattern undergoes significant changes on geometry optimization. This is illustrated for a Type I (*System 9*) and a Type II triple (*System 2*). Inter-base hydrogen bonds present in the (A) crystal structure and (B) optimized geometry of System 9 are shown in broken lines. Magenta color is used to represent the proton mediated hydrogen bond. Similarly inter-base hydrogen bonds present in the (A) crystal structure and (B) optimized geometry of System 2 are also illustrated.

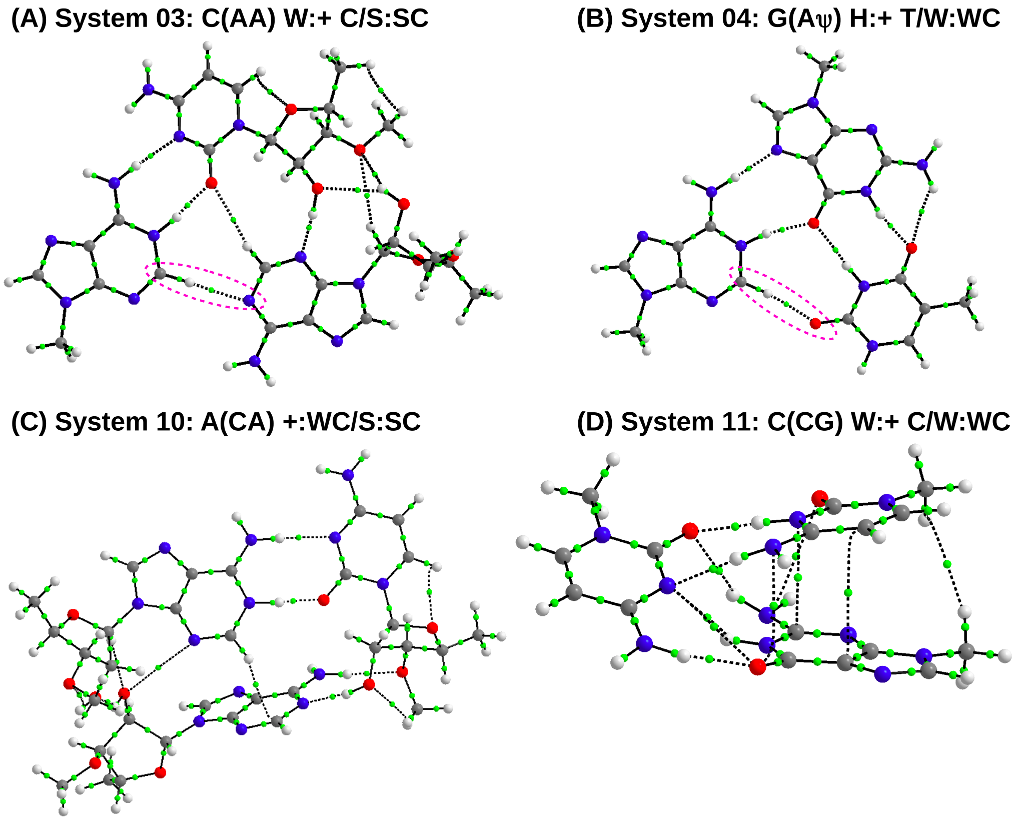


Figure S7: Bond paths obtained from AIM analysis of the four protonated triples are shown. Green spheres represent the bond critical points (BCP). Bond paths corresponding to BCPs having $\nabla^2\rho > 0$ are shown as dotted lines. The C-H$\cdots$N type interaction in *System 3* and C-H$\cdots$O type interaction in *System 4* have been encircled using broken magenta lines.

Table S10: Formation of the *Class I* type protonated base pair changes the charge distribution of the hydrogen bond (HB) donor and acceptor sites of the free edges. q(isolated) represents the partial charges for the isolated nucleobase and q(paired) represents the partial charges after base pair formation. Hydrogen bond donor and acceptor sites are annotated as D and A, respectively. The residues with involve the sugar moiety, have been annotated with 'S' in parenthesis.

| System | Base triple | Protonated base pair | Sites that form HB with the $3^{rd}$ base | | | q (paired) in a.u. | q (isolated) in a.u. | %Change | Change in HB potential |
|---|---|---|---|---|---|---|---|---|---|
| **(1) Type I triples** | | | | | | | | | |
| 1 | A(CC) +:WC/H:WT | A:C +:W Cis | Adenine | N7 | A | -0.465 | -0.489 | -4.9 | -ve |
| | | | | N6 | D | -0.795 | -0.835 | -4.8 | +ve |
| | | | Cytosine | N4 | D | -0.805 | -0.837 | -3.8 | +ve |
| 7 | A(GA) +:HC/S:SC | A:G +:H Cis | Adenine (S) | N3 | A | -0.489 | -0.553 | -11.6 | -ve |
| | | | | O2' | D | -0.761 | -0.771 | -1.3 | +ve |
| | | | | O3' | A | -0.584 | -0.59 | -1.0 | -ve |
| 9 | G(CC) z:WC/W:WC | G:C z:W Cis | Guanine | O6 | A | -0.541 | -0.615 | -12.0 | -ve |
| | | | | N1 | D | -0.657 | -0.669 | -1.8 | +ve |
| | | | | N2 | D | -0.837 | -0.865 | -3.2 | +ve |
| 10 | A(CA) +:WC/S:SC | A:C +:W Cis | Adenine (S) | O2' | A | -0.77 | -0.771 | -0.1 | -ve |
| | | | | N3 | A | -0.475 | -0.553 | -14.1 | -ve |
| | | | Cytosine (S) | O2' | D | -0.774 | -0.786 | -1.5 | +ve |
| | | | | O3' | A | -0.59 | -0.592 | -0.3 | -ve |
| 12 | A(GU) +:HC/S:SC | A:G +:H Cis | Adenine (S) | N3 | A | -0.489 | -0.553 | -11.6 | -ve |
| | | | | O2' | D | -0.761 | -0.771 | -1.3 | +ve |
| 18 | A(CU) +:WC/H:WC | A:C +:W Cis | Adenine | N7 | A | -0.465 | -0.489 | -4.9 | -ve |
| | | | | N6 | D | -0.795 | -0.835 | -4.8 | +ve |
| | | | Cytosine | N4 | D | -0.805 | -0.837 | -3.8 | +ve |
| 20 | G(GU) z:HT/W:WC | G:G z:H Trans | Guanine | O6 | A | -0.543 | -0.615 | -11.7 | -ve |
| | | | | N1 | D | -0.658 | -0.669 | -1.6 | +ve |
| | | | | N2 | D | -0.829 | -0.865 | -4.2 | +ve |
| **(2) Type II triples with CE < 0** | | | | | | | | | |
| 2 | G(CC) H:+T/W:WC | G:C H:+ Cis | Guanine | O6 | A | -0.688 | -0.615 | 11.9 | +ve |
| | | | | N1 | D | -0.645 | -0.669 | -3.6 | +ve |
| | | | | N2 | D | -0.841 | -0.865 | -2.8 | +ve |
| 5 | G(CC) H:+C/W:WC | G:C H:+ Cis | Guanine | O6 | A | -0.686 | -0.615 | 11.5 | +ve |
| | | | | N1 | D | -0.646 | -0.669 | -3.4 | +ve |
| | | | | N2 | D | -0.842 | -0.865 | -2.7 | +ve |
| 6 | G(AC) H:zC/W:WC | G:A H:z Cis | Guanine | O6 | A | -0.669 | -0.615 | 8.8 | +ve |
| | | | | N1 | D | -0.653 | -0.669 | -2.4 | +ve |
| | | | | N2 | D | -0.842 | -0.865 | -2.7 | +ve |
| 8 | G(AC) H:+T/W:WC | G:A H:+ Trans | Guanine | O6 | A | -0.703 | -0.615 | 14.3 | +ve |
| | | | | N1 | D | -0.643 | -0.669 | -3.9 | +ve |
| | | | | N2 | D | -0.842 | -0.865 | -2.7 | +ve |
| 13 | G(CA) W:+C/S:WT | G:C W:+ Cis | Guanine(S) | N2 | D | -0.843 | -0.863 | -2.3 | +ve |
| | | | | O2' | D | -0.762 | -0.77 | -1.0 | +ve |
| 14 | G(CA) W:+C/S:ST | G:C W:+ Cis | Guanine | N2 | D | -0.845 | -0.865 | -2.3 | +ve |
| **(3) Type II triples with CE > 0** | | | | | | | | | |
| 3 | C(AA) W:+C/S:SC | C:A W:+ Cis | Cytosine (S) | O2 | A | -0.733 | -0.693 | 5.8 | +ve |
| | | | | O2' | D | -0.774 | -0.786 | -1.5 | +ve |
| | | | | O3' | A | -0.612 | -0.592 | 3.4 | +ve |
| | | | Adenine | C2 | D | 0.294 | 0.257 | 14.4 | +ve |
| 4 | G(A$\psi$) H:+T/W:WC | G:A H:+ Trans | Guanine | O6 | A | -0.703 | -0.615 | 14.3 | +ve |
| | | | | N1 | D | -0.643 | -0.669 | -3.9 | +ve |
| | | | | N2 | D | -0.842 | -0.865 | -2.7 | +ve |
| | | | Adenine | C2 | D | 0.292 | 0.257 | 13.6 | +ve |

### PDB Ids of the RNA crystal structures studied in this work

#### HD-RNAS (167 files)

URL: http://www.saha.ac.in/biop/www/HD-RNAS.html

2du6, 2zue, 1f7u, 3kfu, 1b23, 2fmt, 1yfg, 4arc, 3vjr, 2bte, 2azx, 3eph, 1o0c, 3knl, 3akz, 1euy, 1zjw, 3amt, 1ffy, 1qu2, 2zni, 3foz, 1evv, 2zm5, 3tup, 1ehz, 3a3a, 3hl2, 3add, 2fk6, 1qf6, 2dr2, 2y10, 2xqd, 3am1, 1c0a, 1il2, 2der, 1gtr, 1n78, 1qtq, 1wz2, 3tvf, 3uye, 4as1, 2ct8, 3knh, 4dh9, 2csx, 2du5, 1h4s, 3rg5, 1ser, 4gaq, 3uz6, 1j1u, 3uz8, 1j2b, 1gax, 1i6u, 2vqe, 3r8n, 3u5b, 2zjr, 3r8s, 1mms, 1vqo, 3v2d, 3u5d, 1un6, 2xg0, 1u8d, 1xok, 2vpl, 1zho, 2nz4, 1hr2, 1gid, 1c9s, 1sj3, 3nkb, 3g78, 2z75, 2r8s, 2gcv, 1m5o, 3cul, 3d2v, 3f2q, 4enc, 2xnz, 3npq, 3irw, 3v7e, 4fe5, 1y26, 3q3z, 3gx5, 3suh, 3dio, 2pxv, 1l9a, 1lng, 3ktw, 1hq1, 2pxb, 2a64, 1u9s, 1p6v, 3r9w, 3r9x, 2b57, 3npb, 1gtn, 1gtf, 3snp, 3pu0, 3pu1, 2xd0, 2zzn, 2zzm, 2ozb, 2gje, 3ptx, 3pu4, 1fir, 3moj, 3hhz, 357d, 3knn, 1s03, 3siv, 2xdb, 2xdd, 2il9, 3nmu, 1kxk, 1vfg, 3pyu, 3kiq, 3kir, 3kit, 3pyn, 3u4m, 3ciy, 1jbr, 3nvk, 3hax, 3ovb, 3ovs, 3p22, 1kh6, 3pla, 1ddy, 3r4f, 3rw6, 2zh1, 3icq, 2gtt, 2nue, 2hvy, 1u6b, 2hw8, 2jea, 3lwv, 1xjr, 3ds7

#### NDB (838 files)

URL: http://rna.bgsu.edu/rna3dhub/nrlist/

1urn, 3iab, 1ooa, 3r4f, 3zp8, 3snp, 4a3e, 4pqv, 1kxk, 4bxx, 3siu, 3la5, 4tzx, 1cvj, 2bh2, 4a3d, 4a3k, 3b5f, 4lx6, 4q5s, 1p6v, 3egz, 364d, 1t0d, 1yfg, 3nvk, 1hys, 4boc, 3hk2, 1i9v, 1mji, 2tra, 1y26, 4n0t, 2du3, 3moj, 2du6, 1jbr, 2d2l, 3cw5, 1h4s, 1qc0, 4ngf, 3nkb, 1f7v, 1fir, 4c4w, 1vfg, 4jf2, 3vjr, 4ol8, 2ake, 4a93, 2dlc, 1vby, 3tup, 1u0b, total 3ouy, 2ozb, 3e2e, 1qu2, 2g3s, 4kr2, 2p7d, 2hoj, 1j1u, 1i6u, 3wqy, 3po2, 1qf6, 1q96, 1b23, 2rfk, 1VY9, 3amu, 2zuf, 4k4w, 4p5j, 3umy, 2hvy, 1eiy, 1qrs, 4mcf, 1VY8, 4qi2, 2zzm, 3cz3, 4lvz, 4db2, 4kr6, 2gjw, 3hjw, 3gtl, 4kr7, 4k4u, 1qbp, 3am1, 3hax, 3m3y, 1h3e, 3qsy, 4aob, 2i82, 4b5r, 1feu, 3dh3, 3al0, 3bnq, 2nok, 1s03, 2pn4, 3gx5, 4kzd, 4ts2,

3fu2, 1d4r, 4l81, 1tfy, 4wkj, 1tfw, 4o26, 4q0b, 3sux, 1nuv, 3p59, 3zla, 1lng, 3htx, 3ciy, 1yyw, 2vpl, 4w92, 1u63, 3hou, 4pqu, 1ykv, 3w3s, 3ucz, 4wfl, 1duq, 4qlm, 4H6F, 1egk, 3a6p, 4jrc, 4qyz, 3f4g, 4qk9, 4kqy, 1qa6, 4g7o, 1zx7, 4csf, 4mgm, 1fjg, 4qk8, 3j7y, 2AW7, 4un5, 4un3, 1un6, 4v19, 4ilm, 3rw6, 1mfq, 3ivn, 3icq, 2o5i, 2oiu, 3slq, 3j7a, 3siv, 1hmh, 4uyk, 4u7u, 3eph, 2quw, 2zzn, 2ho7, 4W23, 1h38, 3U5F, 2der, 2zni, 2zm5, 3pla, 2azx, 1efw, 1il2, 2qbz, 2wj8, 3wfs, 2qux, 1u9s, 1j2b, 2y9h, 2csx, 3d2v, 3q3z, 2qwy, 2d6f, 4k4y, 2cv1, 1gax, 3d0u, 3ol8, 2bte, 4aq7, 2gdi, 3ol7, 4gxy, 4frg, 4rge, 3ol9, 3rg5, 3dir, 3owz, 1et4, 1wz2, 1g1x, 3add, 3adc, 4p9r, 3cun, 3iwn, 3ktw, 4k50, 3p22, 4frn, 2gtt, 4FY3, 2xd0, 4plx, 4uyj, 1m5o, 1u6b, 1VW3, 4eya, 4oo8, 2czj, 4ioa, 4m6d, 3v7e, 1s72, 4QCN, 2il9, 3ivk, 2QBG, 3rkf, 1nbs, 3r1c, 4qjd, 2a64, 3U5H, 3kfu, 1kog, 1VX6, 1gid, 4mgn, 3hga, 3ok4, 1rmv, 4ejt, 3j06, 4lck, 4W21, 1rxa, 4ang, 3ahu, 3pdm, 1vtm, 2x1a, 1mdg, 1laj, 1osu, 2c4z, 2c50, 1p79, 2fz2, 4afy, 4alp, 2xnr, 4lq3, 4jiy, 4n2s, 333d, 3u2e, 2x1f, 1h2c, 4ohy, 4n2q, 3qjp, 2r7u, 2r7v, 4qu6, 4lmz, 4faw, 2r7s, 3o8c, 3mj0, 2tmv, 4r3i, 3hsb, 4ms9, 4ohz, 2r7t, 283d, 3pf4, 3qsu, 3gpq, 3rtj, 2vop, 1g2j, 3b0u, 3nj6, 4dwa, 3q0q, 3q0r, 3q0s, 2a1r, 4jzv, 4jvh, 3k5q, 3k5y, 3k5z, 3k61, 3k62, 3k64, 4tu0, 3pf5, 3rc8, 3t3o, 1b2m, 2voo, 4i67, 2xs7, 4rkv, 3qgc, 1l3z, 1kq2, 4hot, 4hos, 3avt, 4nku, 4lj0, 2asb, 4j7m, 4qu7, 3nna, 2g91, 1n1h, 413d, 3o8r, 3v71, 3v6y, 2xs2, 3bsb, 4j7l, 4oau, 1nb7, 4hor, 1fxl, 2bbv, 1si3, 4nl3, 3nma, 1i5l, 1pgl, 3mij, 3nmr, 4jab, 255d, 2q66, 2r7w, 4oav, 3t5n, 3v74, 3l26, 1uvk, 4rcj, 1g2e, 4kre, 3nnc, 4d25, 1uvi, 3ice, 3gib, 2c0b, 2po1, 4ola, 3pey, 2q1r, 3g9y, 2vnu, 4am3, 3mdg, 3mdi, 4e59, 4u35, 2v7r, 4u34, 259d, 402d, 4cs1, 3iev, 1wpu, 3sqw, 1uvj, 2jea, 3czw, 4fts, 4kxt, 1cwp, 4e78, 2xs5, 4c8y, 4g9z, 3r1e, 4al7, 3nd3, 3q0o, 3i5x, 3i5y, 1wmq, 1av6, 2a8v, 4qm6, 4ii9, 4mdx, 2xgj, 4krf, 4c8z, 2vuq, 2a0p, 3p4c, 3p4d, 4c40, 4f1n, 2xzo, 2b2d, 3jxr, 3oin, 2xli, 1kd5, 438d, 1kfo, 3ie1, 1uvm, 3q0p, 2bx2, 1ddl, 3o7v, 2jlu, 2jlw, 2vrt, 4g0a, 3bx3, 2g4b, 3aev, 3er9, 2von, 1n38, 4gv9, 3kmq, 2f8k, 3knc, 4o8j, 3rer, 1uvl, 1pvo, 4ba2, 4jvy, 4fsj, 2vod, 1zse, 1zdj, 4e58, 2gxb, 3gvn, 421d, 479d, 4msr, 3bnt, 1bmv, 4ht9, 4n48, 3bx2, 4dzs, 4b3g, 4ftb, 3kms, 3nl0, 422d, 472d, 1dqh, 4jrd, 377d, 3ibk, 1m8y, 4ngd, 3qrr, 3bsx, 4nha, 4f3t, 3uld, 3vnu, 1wne, 165d, 2y8y, 439d, 4l8h, 3eqt, 2r22, 2jlz, 1xpo, 1xpr, 1xpu, 1knz, 3fht, 2izn, 2y8w, 3avu, 2v6w, 4fvu, 3l25, 4j39, 3pkm, 4jk0, 4w5n, 3qjj, 3qjl, 1zh5, 3qrp, 1i9x, 1j6s, 4j5v, 2jlx, 3d2s, 2atw, 1n35, 1uvn, 1ytu, 3vnv, 4nlf, 3fs0, 1lnt, 157d, 3cjz, 4jrt, 4knq, 2ix1, 354d, 3avv, 4e6b, 2pjp, 2r20, 433d, 4pei, 4f02, 3bso, 1fuf,

3boy, 2py9, 3m85, 3q2t, 3bsn, 3cgp, 3cgr, 3cgs, 3ssf, 1rna, 4kyy, 2gje, 3dd2, 3m7n, 3nnh, 2ann, 2e9r, 4c9d, 4w5r, 3adi, 1l2x, 4h5p, 1b7f, 2e9z, 387d, 4tv0, 3hxm, 3r2d, 4ay2, 353d, 1csl, 2a43, 466d, 435d, 405d, 3q50, 2nz4, 4pmw, 1f8v, 2izm, 1k8w, 3avw, 1mhk, 397d, 4gha, 2val, 1m8w, 4peh, 3r2c, 4bw0, 1zbi, 1zbl, 4oq8, 3avx, 2bgg, 3ncu, 3lrr, 3k49, 3avy, 2j0s, 7msf, 3ho1, 3tmi, 2xsl, 4pcj, 2w89, 420d, 3og8, 1jid, 1jzv, 4w5t, 2iz9, 4ato, 4kji, 3hm9, 3sn2, 4h8k, 406d, 398d, 1f1t, 1j9h, 3p6y, 4bwm, 2zy6, 1zev, 2jlt, 4m59, 4w5q, 2bu1, 3glp, 2oe5, 4m7d, 4w5o, 1u1y, 2e9t, 3hjf, 1f27, 3lrn, 3tzr, 3sj2, 1o9m, 2ao5, 3r9w, 2db3, 4oo1, 4m7a, 4ill, 1t0k, 4qil, 2ec0, 1e7k, 1sa9, 2hw8, 1saq, 4pdb, 4e5c, 4ijs, 1UTF, 3mqk, 3szx, 1t0e, 4fxd, 4ghl, 2bq5, 5msf, 4jah, 2dr8, 361d, 3syw, 1mwl, 3bnl, 4qqb, 6msf, 3ova, 2fqn, 2az2, 3td0, 4k31, 2grb, 4jng, 3bt7, 1zbh, 1xok, 2o3x, 3td1, 4ktg, 3mei, 2qek, 4bhh, 4u8t, 1br3, 1ec6, 1ntb, 4hkq, 4oog, 2o3v, 4kq0, 409d, 1duh, 1sds, 3rzo, 1yzd, 1q2r, 4gg4, 4nfo, 4qik, 2zi0, 1nlc, 2awe, 2gic, 3ptx, 3pu0, 4iqs, 3s1m, 3s1n, 2f8s, 3bnp, 2zko, 1xpe, 280d, 4j1g, 1gtn, 1gtf, 4o41, 4mce, 1rpu, 2qk9, 4erd, 3erc, 4lgt, 3npn, 3loa, 1yz9, 3ks8, 1xjr, 2nue, 3p4b, 3pu1, 3zc0, 1qcu, 3agv, 4wsb, 3r9x, 4wsa, 4wrt, 3b31, 4py5, 3ftm, 1kh6, 1utd, 3s1r, 3ts0, 1dfu, 429d, 4ig8, 1r9f, 2q1o, 4k27, 3s2d, 1rlg, 4enb, 3wbm, 4oji, 2f8t, 3ts2, 2i91, 3s14, 2ply, 1sdr, 2xb2, 1VW4, 3s49, 3s15, 3ftf, 2nug, 2pxv, 3s16, 1i6h, 3r1d, 3s17, 3e5c, 2b3j, 1qln, 488d, 3nvi, 3nj7, 3po3, 4k4v, 1mzp, 4m30, 3t5q, 4a3g, 4gcw, 1jbs, 4m4o, 1m8v, 2w2h, 1s77, 1yvp, 2ez6, 4kz2, 3fo4, 1msw, 4e48, 3f73, 1r3e, 1f7y, 4c7o, 1e8o, 3bsu, 1ser, 4ifd, 1a9n, 4b3o

#### Larger dataset (2464 files)

URL: https://drive.google.com/open?id=0Bz9N3h6Uh7BYd0hHRmIyUlA3S0E

**X-ray crystal structures with resolution less than 3.5Å (1873 files)**

2xpj, 3hga, 3ok4, 4ejt, 1rxa, 3wzi, 4ang, 3ahu, 2x1a, 1mdg, 1laj, 1osu, 2c4z, 2c50, 2c51, 1p79, 2fz2, 4afy, 4alp, 2xnr, 4qvc, 4lq3, 4jiy, 4n2s, 333d, 3u2e, 2x1f, 1h2c, 1eqq, 4oi0, 4oi1, 4ohy, 4n2q, 3qjp, 2r7u, 2r7v, 4yoe, 4qu6, 4lmz, 2r7s, 3o8c, 3mj0, 4r3i, 4qvd, 3hsb, 4ms9, 4ohz, 2r7r, 2r7t, 283d, 3pf4, 4rcm, 3qsu, 3gpq, 3rtj, 2vop, 1g2j, 3b0u, 3nj6, 4dwa, 3qg9, 3q0q, 3q0r, 3q0s, 3af6, 4s2y, 5bud, 2a1r, 4jzv, 4jvh, 3k5q, 3k5y, 3k5z, 3k61, 3k62, 3k64, 4tu0, 3t3n, 3pf5, 1h2d, 3rc8, 3t3o, 4x9e, 1b2m, 2voo, 4i67, 2xs7, 4rby, 4rbz, 4rc0, 4rkv, 3qgb, 3qgc, 4s2x, 1l3z, 4jzu, 1kq2, 4hot, 4hos, 3avt, 4nku, 4lj0,

2asb, 4j7m, 4qu7, 3nna, 2g91, 4u3p, 4xk0, 1n1h, 4rj1, 413d, 3o8r, 3v71, 5bte, 3v6y, 2xs2, 3bsb, 4j7l, 4oau, 1nb7, 4hor, 1fxl, 2bbv, 1si3, 4nl3, 3nma, 1i5l, 1pgl, 3mij, 3nmr, 4jab, 255d, 2q66, 2r7w, 4oav, 3t5n, 3v74, 3l26, 4tyw, 4tz0, 4tz6, 1uvk, 4rcj, 1g2e, 4kre, 3nnc, 1wrq, 4d26, 4d25, 1uvi, 3ice, 3gib, 2c0b, 2po1, 4ola, 4olb, 3pey, 2q1r, 3g9y, 2vnu, 4am3, 3mdg, 3mdi, 4e59, 4u35, 2v7r, 4u34, 4u37, 259d, 402d, 4u3l, 4u3o, 4u3r, 4u47, 4u78, 4cs1, 3iev, 1wpu, 3sqw, 3sqx, 1uvj, 2jea, 3czw, 4fte, 4fts, 3pew, 4tyy, 4kxt, 1cwp, 4wkr, 4e78, 2xs5, 4c8y, 4g9z, 3r1e, 4al7, 1rxb, 3nd3, 3nd4, 3q0o, 3i61, 3i62, 3i5x, 3i5y, 1wmq, 1av6, 3o3i, 2a8v, 4qm6, 4ii9, 4mdx, 2xgj, 4krf, 4wal, 4c8z, 4u6m, 4u6l, 2vuq, 2a0p, 3p4c, 3p4d, 4c40, 4f1n, 3o6e, 4ht8, 2xzo, 2b2d, 2r7y, 3jxr, 4gv3, 4gv6, 3oin, 4al6, 2xli, 1kd3, 1kd4, 1kd5, 438d, 1kfo, 3ie1, 1uvm, 3q0m, 3q0p, 2bx2, 1ddl, 3o7v, 2jlu, 2jlw, 2vrt, 4g0a, 3bx3, 2g4b, 3aev, 3er9, 2von, 1n38, 4wtk, 4wtl, 4wtm, 4wti, 4wtj, 4gv9, 3kmq, 2xlj, 2f8k, 3knc, 4o8j, 3d0m, 3rer, 1uvl, 5c0y, 1pvo, 4ngg, 4ba2, 4jvy, 4fsj, 2vod, 1zse, 4iqx, 1zdj, 4e58, 2gxb, 3gvn, 421d, 479d, 4msr, 2r7x, 3q0n, 1m8x, 3bnt, 1bmv, 4nh5, 3g0h, 4ht9, 4n48, 3bx2, 4dzs, 4b3g, 4ftb, 3klv, 3kms, 3kna, 3nl0, 422d, 472d, 3jxq, 1efo, 1dqf, 1dqh, 4jrd, 377d, 3ibk, 3q0l, 1m8y, 4ngb, 4ngc, 4ngd, 3qrr, 3bsx, 4nha, 4f3t, 2bs1, 3uld, 3vnu, 1wne, 2g8f, 2g8h, 2g8k, 2g8u, 2g8v, 2g8w, 2gun, 165d, 2y8y, 439d, 4l8h, 3eqt, 2g92, 1g4q, 1fix, 404d, 2ykg, 205d, 2r21, 2r22, 4nh3, 2jly, 2jlz, 1xpo, 1xpr, 1xpu, 1knz, 3fht, 4wta, 4wtc, 4wtf, 4wtg, 2izn, 2iz8, 2c4q, 2c4y, 4wte, 4wtd, 2y8w, 4zld, 3avu, 2v6w, 4fvu, 3l25, 2dqo, 2dqp, 2dqq, 4bpb, 3zd6, 4j39, 1j8g, 3pkm, 4jk0, 2jlv, 4w5n, 3qjj, 3qjl, 4x2b, 1yty, 1zh5, 3qrp, 3adl, 1jb8, 1i9x, 4jgn, 1j6s, 4nh6, 4j5v, 2jlx, 3d2s, 2atw, 3twh, 1n35, 2g8i, 1uvn, 1ytu, 4nmg, 3vnv, 4nxh, 4nlf, 3fs0, 1lnt, 3vyx, 157d, 3cjz, 4jrt, 4knq, 2yjy, 4ed5, 2ix1, 4wzq, 354d, 3avv, 4e6b, 3dw4, 3dw6, 480d, 2pjp, 1pjg, 1pjo, 2r1s, 2r20, 433d, 4pei, 4f02, 2bs0, 4qpx, 3bso, 2ab4, 3zd7, 430d, 1fuf, 3boy, 2py9, 3m85, 3q2t, 3bsn, 3cgp, 3cgq, 3cgr, 3cgs, 3ssf, 437d, 1rna, 4kyy, 3k1v, 2gje, 3dd2, 3m7n, 3nnh, 2ann, 2e9r, 4wzm, 3h5x, 3h5y, 1a34, 4c9d, 4w5r, 3s7c, 3s8u, 3dvz, 3dw5, 3dw7, 483d, 3adi, 1l3d, 1l2x, 3q51, 4h5p, 1b7f, 2e9z, 387d, 4tv0, 3hxm, 5amr, 3r2d, 1q9a, 4ay2, 1msy, 353d, 1csl, 2a43, 2h1m, 464d, 466d, 434d, 435d, 405d, 3q50, 4pmw, 1f8v, 2izm, 1k8w, 3avw, 1mhk, 397d, 4gha, 2val, 1m8w, 4peh, 3r2c, 2anr, 1zl3, 4bw0, 1zbi, 1zdh, 4oq8, 3avx, 2bgg, 3ncu, 3lrr, 4fnj, 3k49, 3k4e, 3avy, 2j0s, 7msf, 4u6k, 1zdi, 3ho1, 3tmi, 2xsl, 4pcj, 1ik5, 2w89, 420d, 3og8, 3iem, 2bny, 1aq3, 2xlk, 1jid, 2oe6, 2oe8, 3gca, 1jzv, 4w5t, 3oij, 2iz9, 4ato, 4kji, 3hm9, 3sn2, 4h8k, 1f1t, 1j9h, 3p6y, 4bwm,

1aq4, 2dr5, 2zy6, 2jlt, 4m59, 4oe1, 4w5q, 2b2e, 3gm7, 2bu1, 3glp, 2dr7, 2oe5, 4m7d, 4w5o, 2b2g, 1u1y, 2e9t, 3hjf, 1f27, 2dr9, 2zh1, 2zh7, 2zh9, 3lrn, 3tzr, 4wcq, 4wcs, 3sj2, 1o9m, 2ao5, 3r9w, 2db3, 4oo1, 2j0q, 4m7a, 4ill, 1t0k, 2drb, 2zh2, 2zh3, 2zh4, 2zh5, 2zh8, 2zhb, 4qil, 4pgy, 2ec0, 1e7k, 1sa9, 2hw8, 4f8v, 4wcr, 1saq, 4pdb, 4e5c, 4ijs, 3mqk, 2xdb, 3szx, 4j50, 4yn6, 1t0e, 4p20, 4fxd, 4k32, 4ghl, 3ex7, 2bq5, 5msf, 4jah, 2dr8, 299d, 361d, 3syw, 301d, 1j7t, 4gpx, 1lc4, 1mwl, 3dvv, 3bnl, 4qqb, 6msf, 5amq, 2dvi, 2dra, 2zha, 3ova, 1q29, 2fqn, 2az0, 2az2, 2g5k, 3td0, 4k31, 2grb, 4jng, 2hyi, 3bt7, 1zbh, 2zh6, 1xok, 1zdk, 2f4s, 2f4u, 2o3x, 4gpw, 3td1, 3s4p, 2be0, 2bee, 4gpy, 2pwt, 4ktg, 3mei, 3c44, 2qek, 4bhh, 4wan, 4u8t, 1br3, 1ec6, 1di2, 4jnx, 300d, 359d, 1nta, 1ntb, 4hkq, 1nyi, 4oog, 3wru, 1z7f, 4pmi, 2et4, 2et5, 2o3w, 2et3, 2et8, 2o3v, 2o3y, 1yrj, 4kq0, 1rc7, 2oiy, 2oj0, 4lg2, 409d, 1duh, 1sds, 3rzo, 1yzd, 1z79, 4nfp, 4nfq, 1q2r, 4gg4, 4nfo, 4qik, 2esj, 2zi0, 3far, 1nlc, 1y90, 2awe, 2gic, 3ptx, 3pu0, 3pu4, 379d, 4iqs, 3s1m, 3s1n, 3rzd, 2f8s, 4f8u, 4pdq, 3bnp, 2zko, 5bs3, 2b8r, 1xpf, 1xp7, 1xpe, 280d, 4j1g, 1gtn, 1gtf, 4phy, 2esi, 4o41, 4mce, 1rpu, 2qk9, 2fcx, 2fcy, 2fd0, 4erd, 1o3z, 3erc, 4lgt, 3mj3, 3mja, 3mjb, 3npn, 4en5, 3loa, 1ze2, 2oij, 1yz9, 3ks8, 4rwp, 3vrs, 4s3n, 1xjr, 4enc, 2nue, 1y6s, 1wvd, 1y6t, 1y73, 1y95, 1y99, 4rne, 3p4b, 3pu1, 3zc0, 1qcu, 4r8i, 3agv, 4wsb, 3r9x, 4wsa, 4wrt, 3b31, 4py5, 4zcf, 3lqx, 2b8s, 1y3o, 1y3s, 1yxp, 3ftm, 1kh6, 1utd, 3s1r, 1yy0, 1q2s, 3ts0, 1dfu, 429d, 2pn3, 4ig8, 1r9f, 2q1o, 4k27, 3s2h, 3s2d, 2pxl, 4ena, 1rlg, 4enb, 3wbm, 4oji, 2f8t, 1yyk, 3hhz, 3ts2, 2i91, 3s14, 2uwm, 3fte, 2ply, 462d, 1sdr, 1yyo, 2xb2, 4a3b, 3s49, 3s15, 3ftf, 2pxb, 2pxe, 2pxu, 2nug, 2pxd, 2pxf, 2pxk, 2pxp, 2pxq, 2pxt, 2pxv, 1hq1, 4tux, 4rwn, 4rwo, 3s16, 3vyy, 1i6h, 4enn, 2f4t, 3s1q, 3r1d, 2qkb, 4p97, 3s17, 1dul, 3e5e, 3e5f, 3e5c, 1zbl, 4qoz, 4l8r, 2b3j, 1qln, 2fk6, 4u38, 4msb, 488d, 3nvi, 4m2z, 3nj7, 4v4f, 1c9s, 3po3, 1mzp, 4m30, 2nuf, 3t5q, 4a3g, 4gcw, 1jbs, 4m4o, 1m8v, 2w2h, 1s77, 1yvp, 3hvr, 1jbt, 4xw7, 4xwf, 2ez6, 4kz2, 3fo4, 3ok2, 1msw, 4e48, 3f73, 1r3e, 4q5v, 4tuw, 1f7y, 4c7o, 1e8o, 1kuq, 3bsu, 1zz5, 2a04, 1ser, 3gog, 4ifd, 1s76, 1a9n, 1dk1, 4b3o, 3gs5, 5c7w, 3gao, 2xnz, 3fo6, 4fe5, 3g4m, 3ger, 1urn, 3iab, 4wcp, 1ooa, 3r4f, 2b57, 3ges, 3got, 2ees, 2eet, 2eeu, 2eev, 2eew, 3hov, 4a3c, 5c7u, 4fej, 4fel, 2g9c, 4fen, 4feo, 4fep, 3snp, 4a3e, 4pqv, 2oeu, 1kxk, 4bxx, 3siu, 4g6s, 4g6p, 3bbm, 1vtq, 3b5a, 3b5s, 3la5, 4tzx, 4tzy, 1cvj, 2bh2, 3hoy, 4a3d, 4a3k, 4ycp, 3b5f, 3b91, 3bbk, 3b58, 2oue, 4lx6, 4lx5, 4q5s, 4x4v, 1p6v, 3egz, 357d, 364d, 1t0d, 3l0u, 1yfg, 4tna, 4xnr, 3nvk, 1hys, 4boc, 4a3l, 1wsu, 1ehz, 3tra, 3hk2, 6tna, 1tn2, 1i9v, 3bbi, 1mji, 2tra, 1y27,

1y26, 4n0t, 2det, 2du3, 2du4, 4g6r, 1x9c, 1tn1, 1zev, 2hop, 1zft, 1vc5, 1tra, 3moj, 1gsg, 2nqp, 2du6, 2du5, 1jbr, 4arc, 4pkd, 2d2l, 4a3f, 3cw5, 3cw6, 4tra, 1gts, 1h4s, 1qc0, 1evv, 4ngf, 2nvt, 4x4s, 2d2k, 1f7v, 1zfv, 2bcy, 2fgp, 3gs8, 2p7f, 1fir, 4c4w, 4x4q, 2npy, 1x9k, 1vfg, 1qtq, 1zfx, 1vc6, 4qei, 4jf2, 3vjr, 4ol8, 2ake, 4a93, 2dlc, 4nyg, 3ovs, 3ov7, 1zjw, 2oih, 2oj3, 2bcz, 1vbx, 1vby, 1vc0, 3tup, 1u0b, 1vbz, 1h4q, 2hom, 4yvi, 4yvj, 4yvk, 4x0b, 4wc3, 3ouy, 2npz, 2rd2, 1euq, 3gs1, 1f7u, 4x4o, 1exd, 2ozb, 1sjf, 1sj3, 1sj4, 3e2e, 2hok, 1qu3, 4nyb, 4nyc, 1ffy, 1qu2, 4nyd, 2g3s, 4jxx, 1euy, 4kr2, 4jyz, 4jxz, 1gtr, 2re8, 4pr6, 2hoj, 2hol, 1j1u, 1i6u, 2bj6, 3x1l, 4wc4, 2dr2, 4wc5, 4x4r, 3ovb, 3wqz, 1o0c, 3wqy, 2hoo, 4wj4, 4zc7, 1q93, 4wc6, 4wc2, 4wc7, 3po2, 4k4t, 3bnn, 4prf, 1ob2, 4kr3, 4by7, 3bno, 1qf6, 2vum, 1q96, 3a3a, 4xjn, 1b23, 2rfk, 1o0b, 1c0a, 4k4s, 3amt, 3amu, 4kr9, 2zuf, 4k4w, 1mme, 1qrt, 4ari, 4p5j, 3u56, 3umy, 2hvy, 1eiy, 1drz, 4rdx, 1qru, 2zue, 1qrs, 5btm, 4mcf, 4qi2, 2zzm, 3lwv, 3cz3, 4lvw, 4lvx, 4lvy, 4lvz, 4db2, 2e2j, 4kr6, 3lwr, 2gjw, 3hjw, 4as1, 4qvi, 3u4m, 4qg3, 4lw0, 3gtj, 2iy5, 3gtl, 4kr7, 4k4u, 3lwq, 3lwo, 3lwp, 1qbp, 3sd3, 4x4u, 3am1, 3hax, 4lvv, 4q9q, 4q9r, 2yu9, 2e2i, 2nvq, 1r9t, 4y52, 4y7n, 3m3y, 1h3e, 3qsy, 1k9w, 4aob, 3iqp, 2gis, 2i82, 3iqn, 2ydh, 4b5r, 3iqr, 3zju, 1feu, 3dh3, 3al0, 4rum, 3bnq, 2nok, 1s03, 2pn4, 3gx6, 3gx7, 4kze, 3gx2, 2fcz, 3gx3, 3gx5, 3hl2, 4x4n, 4kzd, 4ts2, 3fu2, 1d4r, 4yb1, 4yhw, 1zci, 4l81, 4oqu, 3zjt, 1tfy, 3bns, 4wkj, 1tfw, 4o26, 4puo, 4ts0, 4q0b, 3suh, 3sux, 3suy, 1nuj, 1nuv, 3p59, 1z43, 3zla, 3irw, 3muv, 4rzd, 1lng, 3htx, 3ciy, 1yyw, 3mut, 3zjv, 3mxh, 3mum, 3mur, 3bnr, 2vpl, 4w92, 4pwd, 4x4t, 1u63, 3hou, 4pqu, 1ykq, 1ykv, 3ucu, 4w90, 3w3s, 1yls, 3ucz, 3ud4, 3ud3, 4wfl, 1duq, 4qlm, 3f30, 3f2t, 3f2x, 3f2y, 3f2w, 3f2q, 3nmu, 2xdd, 1egk, 3a6p, 4jrc, 4qyz, 3f4h, 3f4e, 3f4g, 4qk9, 2yif, 4kqy, 4tvx, 4qln, 1qa6, 4g7o, 1zx7, 2yie, 5btp, 4qka, 4csf, 4mgm, 1y39, 4qk8, 1hc8, 5axw, 5czz, 1mms, 4x4p, 4un5, 4un3, 4un4, 1un6, 4ilm, 3trz, 3rw6, 3skr, 1l9a, 3ds7, 3skw, 3skl, 3skt, 4p3e, 1mfq, 4znp, 3ivn, 3icq, 2o5i, 4k4x, 3slm, 2oiu, 3slq, 3skz, 406d, 2qus, 4meg, 1ddy, 3siv, 3ski, 1hmh, 3b4c, 2o5j, 2ppb, 4uyk, 4u7u, 3epk, 3epj, 3ndb, 3b4a, 2ho6, 2gcv, 2h0w, 5cd4, 3eph, 2quw, 2zzn, 4meh, 3b4b, 2ho7, 2gcs, 2h0x, 4zt0, 1h38, 4zt9, 2der, 2zni, 4k4z, 2zxu, 2zm5, 3pla, 3npq, 2azx, 1efw, 1il2, 2qbz, 2wj8, 3wfs, 2qux, 3foz, 2deu, 1u9s, 1j2b, 2y9h, 1asy, 2cky, 2csx, 2ct8, 1g59, 3d2x, 3d2g, 3d2v, 2cv0, 3q3z, 2qwy, 2d6f, 1asz, 1zho, 2byt, 2cv1, 2cv2, 1n78, 1gax, 1ivs, 2v0g, 4erj, 4erl, 3d0x, 3d0u, 2fmt, 3ol8, 1s0v, 2bte, 2dxi, 1n77, 3ol6, 4aq7, 2gdi, 4nya, 3ol7, 4gxy, 4frg, 2r8s, 4rgf, 4rge, 3olb, 3ol9, 4cqn, 4yb0, 3rg5, 4yaz, 3zgz, 3dig,

3dis, 3dj0, 3dil, 3diz, 3diq, 3dix, 3dim, 3dir, 3dj2, 3dio, 3diy, 3oxj, 3owz, 1et4, 1wz2, 2qkk, 3oxb, 3oxd, 3oww, 3owi, 3ox0, 3oxe, 3oxm, 3k0j, 1g1x, 4y1m, 3add, 3adc, 3adb, 4p9r, 3cun, 3iwn, 4p95, 3ktw, 4k50, 3cul, 4xco, 3p22, 4frn, 2v3c, 4y1i, 4y1j, 4wfm, 2gtt, 4v9e, 2xd0, 4yco, 4plx, 4uyj, 1m5o, 1ttt, 1m5k, 1ob5, 1u6b, 3bo3, 1m5v, 1m5p, 4eya, 1zzn, 3bo2, 4v99, 4oo8, 2czj, 4m6d, 3v7e, 2il9, 3ivk, 3rkf, 4pjo, 1nbs, 3hhn, 3r1l, 3r1h, 3r1c, 4qjd, 2a64, 3akz, 3wfr, 1l8v, 3kfu, 1kog, 1hr2, 1gid, 3pdr, 4mgn, 4oq9, 4nia, 3bwp, 4lck, 3eoh, 4fax, 4fau, 4e8p, 4e8r, 4e8q, 4e8n, 4fb0, 4e8m, 3eog, 4faq, 4e8k, 3igi, 4faw, 3g78, 4far, 4e8t, 1fg0, 3g9c, 3g96, 3g8s, 3g8t, 3l3c, 2nz4, 4v8i, 4v8h, 4v7x, 4v7z, 4v7y, 4v7w, 4v8g, 1fka, 4v6c, 4v7u, 4v7v, 4v7s, 4v7t, 2uxd, 4u1u, 4u1v, 4u20, 4u25, 4u27, 4u26, 4u24, 4v84, 4v85, 4wf1, 4v83, 2hhh, 2zm6, 4yhh, 4v95, 4lf6, 4ji3, 4dv7, 4dr3, 4khp, 4lf9, 1ibk, 4lfb, 4ji1, 4lf4, 1hr0, 4lf7, 4lf8, 1i94, 4duy, 4dr2, 4dv6, 4ji0, 1hnz, 1hnw, 1hnx, 1fjg, 1j5e, 4ji7, 1xnq, 2vqe, 4gkk, 2uxb, 1xmo, 2vqf, 1xnr, 1xmq, 4gkj, 4k0k, 4jv5, 2e5l, 2uxc, 1n33, 1ibm, 2uub, 2uuc, 3t1h, 3t1y, 4b3s, 2uua, 2uu9, 4aqy, 4b3r, 4jya, 4b3t, 4dr5, 1ibl, 1n32, 4b3m, 4v6a, 4v9s, 4v9r, 4dr6, 4v9a, 1vy6, 4v9n, 1vy7, 4v90, 4v9h, 1vvj, 4l47, 4lsk, 4lnt, 4lt8, 4tue, 4w2h, 4v5a, 4v9k, 4v9l, 4woi, 4v9c, 4v64, 4v4h, 4v4q, 4v54, 4v52, 4v57, 4v5e, 4v9b, 4v5j, 4v8x, 4v67, 4v50, 4v63, 4v7j, 4wu1, 4v5k, 4wzd, 4v51, 4v6g, 4v7m, 4wzo, 4v7l, 1vy4, 4wro, 1vy5, 4w2i, 4wsd, 4wt1, 4v9i, 4wqr, 4w2g, 4w2f, 4v5d, 4v5c, 4wq1, 4v5l, 4u4z, 4u3m, 4u53, 4u4y, 4u4q, 4u56, 4u3n, 4u55, 4u4n, 4u51, 4u4u, 4u52, 4u50, 4u4r, 4u3u, 4v8n, 4v5s, 4v5q, 4v8b, 4v5p, 4v5r, 4wr6, 4v8d, 4wsm, 4wra, 4u6f, 4v88, 1p9x, 2o44, 2ogm, 1ond, 2aar, 1jzx, 1j5a, 1k01, 1jzy, 3pip, 1njp, 3pio, 2d3o, 3dll, 2zjq, 3cf5, 2zjr, 4io9, 4ioa, 4z8c, 1ffk, 1y69, 2qa4, 3g4s, 3ccj, 1w2b, 2qex, 1n8r, 3ow2, 1yjn, 3ccr, 1yjw, 1yj9, 3g6e, 1yi2, 2otl, 1yit, 3i56, 1nji, 3ccs, 1k73, 2otj, 1s72, 1yij, 3cc2, 1jj2, 3cc7, 3ccl, 3ccq, 3cc4, 3cce, 3ccv, 3ccu, 3ccm, 1kc8, 1k8a, 1k9m, 1q81, 1q82, 1kd1, 1m1k, 1m90, 1vqo, 3g71, 1vqk, 1yhq, 3cpw, 1vq8, 1vq9, 3cd6, 1vq6, 3cme, 3cxc, 1kqs, 1vq4, 1vq5, 1q86, 3cma, 1vqn, 1vq7, 1vql, 1vqp, 1vqm, 1xbp, 1sm1, 1nwx, 1nkw, 1qvg, 1nwy, 1qvf, 4v9f, 4v8a, 4zer, 4v9q, 4v9p, 4v9o, 4wqy, 4www, 4wqf, 4y4p, 4y4o, 4wqu, 4wpo, 4z3q, 4z3s, 4z3r, 4v8q, 4wt8, 4v6f, 4v8c, 4v87, 4v8f, 4v8e, 4w2e, 4ybb, 4v9d

**Solution NMR structures (591 files)**

124d, 176d, 17ra, 1a1t, 1a3m, 1a4d, 1a4t, 1a51, 1a60, 1a9l, 1ac3, 1afx, 1ajf, 1ajl, 1ajt, 1aju, 1akx,

1al5, 1am0, 1anr, 1aqo, 1arj, 1ato, 1atv, 1atw, 1aud, 1b36, 1bau, 1bgz, 1biv, 1bj2, 1bn0, 1bvj, 1byj, 1bz2, 1bz3, 1bzt, 1bzu, 1c0o, 1c2q, 1c4l, 1cq5, 1cql, 1cx5, 1d0t, 1d0u, 1d6k, 1drr, 1dz5, 1e4p, 1e95, 1ebq, 1ebr, 1ebs, 1efs, 1eht, 1ei2, 1ejz, 1eka, 1ekd, 1ekz, 1elh, 1esh, 1esy, 1etf, 1etg, 1exy, 1f5g, 1f5h, 1f5u, 1f6u, 1f6x, 1f6z, 1f78, 1f79, 1f7f, 1f7g, 1f7h, 1f7i, 1f84, 1f85, 1f9l, 1feq, 1fhk, 1fje, 1fl8, 1fmn, 1fnx, 1fqz, 1fyo, 1fyp, 1g3a, 1g70, 1guc, 1h0q, 1hg9, 1hji, 1hlx, 1hs1, 1hs2, 1hs3, 1hs4, 1hs8, 1hwq, 1i3x, 1i3y, 1i46, 1i4b, 1i4c, 1i9f, 1i9k, 1idv, 1ie1, 1ie2, 1ik1, 1ikd, 1j4y, 1jo7, 1jox, 1jp0, 1jtj, 1jtw, 1ju1, 1ju7, 1jur, 1jwc, 1jzc, 1k1g, 1k2g, 1k4a, 1k4b, 1k5i, 1k6g, 1k6h, 1k8s, 1kaj, 1kis, 1kka, 1kks, 1koc, 1kod, 1kos, 1kp7, 1kpd, 1kpy, 1kpz, 1l1c, 1l1w, 1lc6, 1ldz, 1lmv, 1lpw, 1luu, 1lux, 1lvj, 1m5l, 1m82, 1me1, 1mfj, 1mfk, 1mfy, 1mis, 1mnb, 1mnx, 1mt4, 1muv, 1mv1, 1mv2, 1mv6, 1mwg, 1my9, 1n53, 1n66, 1n8x, 1na2, 1nbk, 1nbr, 1nc0, 1nem, 1ntq, 1nts, 1ntt, 1nxr, 1nyb, 1nz1, 1o15, 1okf, 1oln, 1oo7, 1oq0, 1osw, 1ow9, 1p5m, 1p5n, 1p5o, 1p5p, 1pbl, 1pbm, 1pbr, 1pjy, 1q75, 1q8n, 1qc8, 1qd3, 1qes, 1qet, 1qfq, 1qwa, 1qwb, 1r2p, 1r3x, 1r4h, 1r7w, 1r7z, 1rau, 1raw, 1rfr, 1rgo, 1rht, 1rkj, 1rng, 1rnk, 1roq, 1rrd, 1rrr, 1s2f, 1s34, 1s9l, 1s9s, 1scl, 1slo, 1slp, 1sy4, 1syz, 1szy, 1t28, 1t2r, 1t4l, 1t4x, 1tbk, 1tfn, 1tjz, 1tlr, 1tob, 1tut, 1txs, 1u2a, 1u3k, 1u6p, 1ull, 1uts, 1uud, 1uui, 1uuu, 1vop, 1wks, 1wts, 1wtt, 1wwd, 1wwe, 1wwf, 1wwg, 1xhp, 1xsg, 1xsh, 1xst, 1xsu, 1xv0, 1xv6, 1xwp, 1xwu, 1yfv, 1yg3, 1yg4, 1ylg, 1ymo, 1yn1, 1yn2, 1ync, 1yne, 1yng, 1ysv, 1z2j, 1z30, 1z31, 1zbn, 1zc5, 1zif, 1zig, 1zih, 219d, 28sp, 28sr, 2a9l, 2a9x, 2ad9, 2adb, 2adc, 2adt, 2aht, 2ap0, 2ap5, 2au4, 2awq, 2b6g, 2b7g, 2bj2, 2c06, 2cd1, 2cd3, 2cd5, 2cd6, 2cjk, 2d17, 2d18, 2d19, 2d1a, 2d1b, 2dd1, 2dd2, 2dd3, 2err, 2es5, 2ese, 2euy, 2evy, 2f4x, 2f87, 2f88, 2fdt, 2fey, 2fy1, 2g1g, 2g1w, 2gbh, 2gio, 2gip, 2gm0, 2grw, 2gv3, 2gv4, 2gvo, 2h49, 2hem, 2hgh, 2hns, 2hua, 2i2y, 2i7e, 2i7z, 2ihx, 2irn, 2iro, 2ixy, 2ixz, 2jpp, 2jq7, 2jr4, 2jrg, 2jrq, 2jse, 2jsg, 2jtp, 2juk, 2jwv, 2jxq, 2jxs, 2jxv, 2jyf, 2jyh, 2jyj, 2jym, 2k3z, 2k41, 2k4c, 2k5z, 2k65, 2k66, 2k7e, 2k95, 2k96, 2kbp, 2kd4, 2kd8, 2kdq, 2ke6, 2kez, 2kf0, 2kfy, 2kg0, 2kg1, 2kgp, 2kh9, 2khy, 2km8, 2kmj, 2koc, 2kp3, 2kp4, 2kpc, 2kpd, 2kpv, 2krl, 2krp, 2krq, 2krv, 2krw, 2kry, 2krz, 2ktz, 2ku0, 2kur, 2kuu, 2kuv, 2kuw, 2kvn, 2kwg, 2kx5, 2kx8, 2kxm, 2kxn, 2kxz, 2ky0, 2ky1, 2ky2, 2kyd, 2kye, 2kzl, 2l1f, 2l1v, 2l2j, 2l2k, 2l3c, 2l3e, 2l3j, 2l41, 2l5d, 2l5z, 2l6i, 2l8c, 2l8f, 2l8h, 2l8u, 2l8w, 2l94, 2l9e, 2la5, 2la9, 2lac, 2lar, 2lb4, 2lbj, 2lbk, 2lbl, 2lbq, 2lbr, 2lbs, 2lc8, 2ldl, 2ldt, 2ldz, 2leb, 2lec, 2lhp, 2li4, 2li8, 2ljj, 2lk3, 2lkr, 2lp9, 2lpa, 2lps, 2lpt, 2lqz, 2lu0,

2lub, 2lun, 2lup, 2lv0, 2lvy, 2lwk, 2lx1, 2m12, 2m18, 2m1o, 2m1v, 2m21, 2m22, 2m23, 2m24, 2m39, 2m4q, 2m4w, 2m57, 2m58, 2m5u, 2m8d, 2m8k, 2mb0, 2meq, 2mer, 2mf0, 2mf1, 2mfc, 2mfd, 2mfe, 2mff, 2mfg, 2mfh, 2mgz, 2mhi, 2mi0, 2mis, 2miy, 2mjh, 2mki, 2mkk, 2mkn, 2mn0, 2mnc, 2mqt, 2mqv, 2ms0, 2ms1, 2ms5, 2mtj, 2mtk, 2mtv, 2mvs, 2mxj, 2mxk, 2mxl, 2mxy, 2mz1, 2n1q, 2n2o, 2n2p, 2n3q, 2n3r, 2n4j, 2o32, 2o33, 2o81, 2o83, 2oj7, 2oj8, 2oom, 2p89, 2pcv, 2pcw, 2pn9, 2qh2, 2qh3, 2qh4, 2rlu, 2rn1, 2ro2, 2rp0, 2rp1, 2rpk, 2rpt, 2rqc, 2rqj, 2rra, 2rrc, 2rs2, 2rsk, 2ru3, 2ru7, 2tob, 2tpk, 2u2a, 2xc7, 2xeb, 2xfm, 2y95, 2yh1, 3php, 484d, 4a4r, 4a4s, 4a4t, 4a4u, 4b8t, 4bs2, 4by9, 4cio, 5a17, 5a18, 8drh, 8psh

### References


(1) Becke, A. D. Density-functional exchange-energy approximation with correct asymptotic behavior. *Phys. Rev. A* **1988**, *38*, 3098–3100.

(2) Lee, C.; Yang, W.; Parr, R. G. Development of the Colle-Salvetti correlation-energy formula into a functional of the electron density. *Phys. Rev. B* **1988**, *37*, 785–789.

(3) Miehlich, B.; Savin, A.; Stoll, H.; Preuss, H. Results obtained with the correlation energy density functionals of Becke and Lee, Yang and Parr. *Chem. Phys. Lett.* **1989**, *157*, 200–206.

(4) Yanai, T.; Tew, D. P.; Handy, N. C. A new hybrid exchange–correlation functional using the Coulomb-attenuating method (CAM-B3LYP). *Chem. Phys. Lett.* **2004**, *393*, 51–57.

(5) Grimme, S.; Antony, J.; Ehrlich, S.; Krieg, H. A consistent and accurate ab initio parametrization of density functional dispersion correction (DFT-D) for the 94 elements H-Pu. *J. Chem. Phys.* **2010**, *132*, 154104.

(6) Grimme, S.; Ehrlich, S.; Goerigk, L. Effect of the damping function in dispersion corrected density functional theory. *J. Comput. Chem.* **2011**, *32*, 1456–1465.

(7) Adamo, C.; Barone, V. Toward reliable density functional methods without adjustable parameters: The PBE0 model. *J. Chem. Phys.* **1999**, *110*, 6158–6170.

(8) Chai, J.-D.; Head-Gordon, M. Long-range corrected hybrid density functionals with damped atom–atom dispersion corrections. *Phys. Chem. Chem. Phys.* **2008**, *10*, 6615–6620.

(9) Antony, J.; Brüske, B.; Grimme, S. Cooperativity in noncovalent interactions of biologically relevant molecules. *Phys. Chem. Chem. Phys.* **2009**, *11*, 8440–8447.
